# Functional, genomic, and transcriptomic insights into Linear Low-Density Polyethylene (LLDPE) biodegradation by landfill-derived *Brucella intermedia*

**DOI:** 10.1101/2025.11.09.687417

**Authors:** Ifthikhar Zaman, Mahdi Muhammad Moosa, M. Mahboob Hossain

**Affiliations:** Department of Biotechnology, BRAC University, Dhaka 1212, Bangladesh; Department of Microbiology, BRAC University, Dhaka 1212, Bangladesh

**Author notes:** Corresponding Author (MMM). These authors contributed equally to this work.

**Keywords:** LLDPE biodegradation, plastic-degrading bacteria, environmental biotechnology, whole-genome sequencing, environmental adaptation

## Abstract

The global accumulation of linear low-density polyethylene (LLDPE) waste has created an urgent need for sustainable biodegradation strategies. Here, we report the identification and characterization of two landfill-derived *Brucella intermedia* isolates associated with LLDPE biodegradation. Soil samples collected from the Matuail landfill in Dhaka, Bangladesh, were screened on minimal salt medium supplemented with LLDPE. Both isolates demonstrated sustained growth under carbon-limited conditions in which LLDPE was the principal carbon source. LLDPE recovered from bacterial cultures showed chemical and surface morphological changes, as supported by Fourier-transform infrared (FTIR) spectroscopy, droplet-spreading analysis, and scanning electron microscopy (SEM). FTIR analysis revealed additional carbonyl, hydroxyl, and C-O bands consistent with oxidative modification of the polymer surface, while SEM showed roughening, fissures, and fragmentation-like surface features. Whole-genome sequencing identified both isolates as closely related but distinct strains of *Brucella intermedia* and revealed enrichment of candidate functions associated with oxidative activation, depolymerization, and downstream assimilation. Transcriptomic analysis under LLDPE growth further showed expression of multiple candidate plastic degradation- and biofilm-associated genes. Resistome, virulome, and phenotypic antimicrobial susceptibility analyses indicated that both isolates are environmentally resilient, low-pathogenic variants with a limited intrinsic resistome. Both strains also exhibited biofilm-associated growth under hydrocarbon-substituted and plastic-associated conditions. Collectively, these findings identify environmental *B. intermedia* as an unexpected plastic-associated lineage and expand current understanding of the ecological and genomic diversity of bacteria linked to LLDPE biodegradation.

**Environmental Implication:** This study identifies landfill-derived *Brucella intermedia* strains that can support biodegradation of linear low-density polyethylene (LLDPE), a major persistent plastic pollutant in landfill environments. Functional, genomic, and transcriptomic evidence indicate that these isolates use coordinated oxidative, depolymerization, and biofilm-associated responses during growth under plastic-associated, carbon-limited conditions. Their low-pathogenic environmental profiles and limited intrinsic resistomes further support their relevance as candidates for future biodegradation-focused biotechnological research. Overall, these findings expand the known diversity of plastic-associated bacteria and highlight naturally adapted landfill microorganisms as promising resources for sustainable strategies to mitigate plastic accumulation in terrestrial ecosystems.

**Graphical Abstract:** 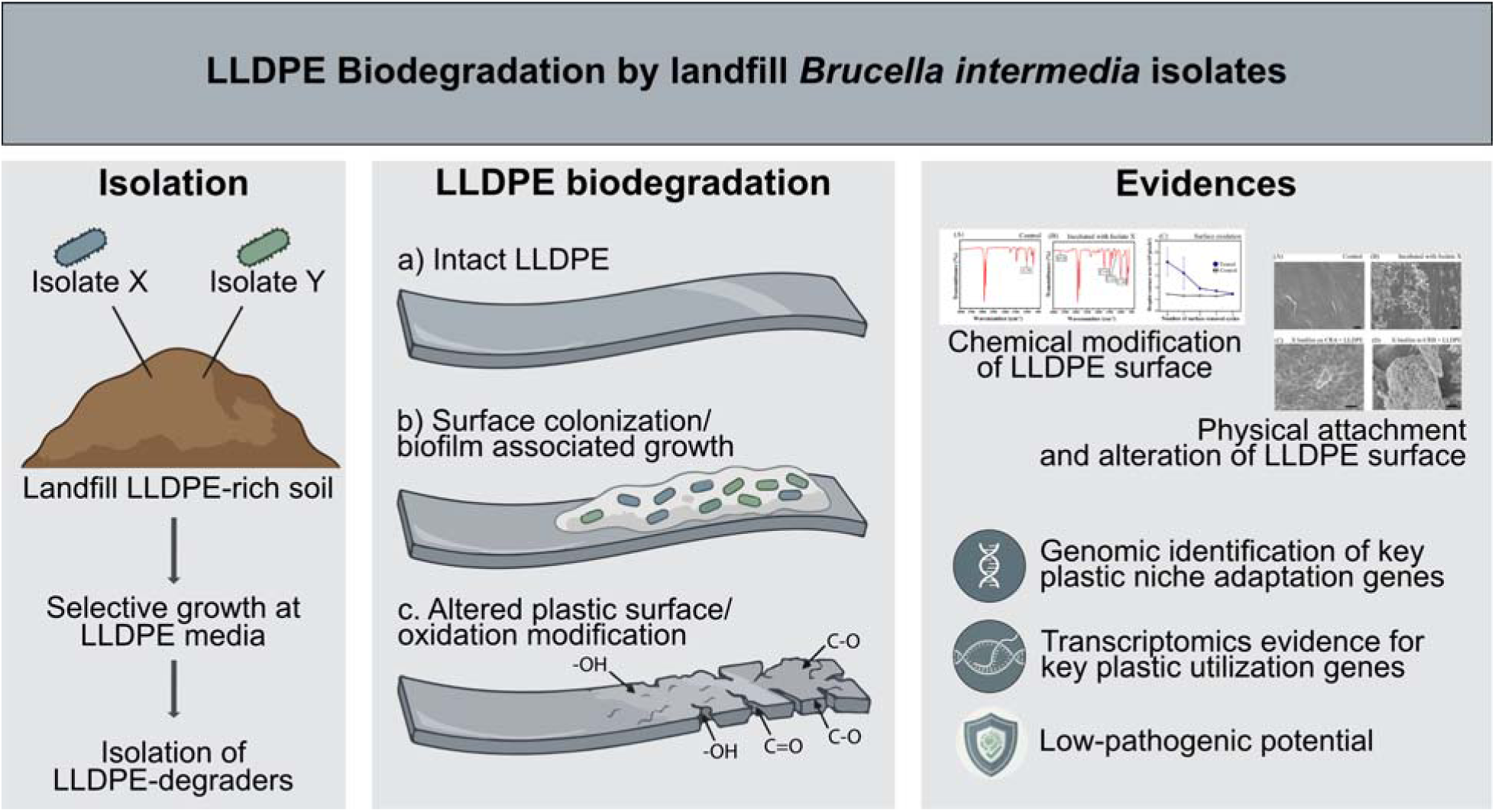

## 1. Introduction

The global plastic pollution crisis has driven significant research into microbial systems capable of degrading recalcitrant polymers, especially polyethylene (PE). As non-biodegradable plastics, PE polymers can persist for centuries and enter food chains through contaminated microplastics (Chamas et al., 2020). Recent studies have confirmed the presence of microplastics in human blood, lungs, and placenta, highlighting potential health risks (Leslie et al., 2022; Ragusa et al., 2021). Among all plastic types, the environmental accumulation of PE waste remains a critical global concern due to its high resistance to degradation and widespread use (Gorish et al., 2024). PE’s chemical inertness, attributed to the strong C–C and C–H bonds, renders its natural removal from the environment extremely slow.

A variety of ethylene-derived polymers are commercially available, with different physical properties and end utilities, including High-Density Polyethylene (HDPE), Low-Density Polyethylene (LDPE), Linear Low-Density Polyethylene (LLDPE), and Polyethylene Terephthalate (PET). Among these, LLDPE is extensively used in food packaging, agricultural films, and household items (Gorish et al., 2024) and therefore accumulates widely in the environment. In this study, we focused on LLDPE, one of the most widely utilized PE types globally (Peacock, 2000; Rahimi and García, 2017). LLDPE is a copolymer primarily composed of ethylene as the main monomer, with small amounts of α-olefins (such as 1-butene, 1-hexene, or 1-octene) as comonomers. These comonomers introduce short-chain branching along the polyethylene backbone, providing distinct properties compared to HDPE. Structurally, LLDPE consists mainly of saturated hydrocarbon chains (–CH –CH – units), but due to comonomer incorporation, it is not a simple linear polymer of ethylene. The introduction of short branches decreases crystallinity and increases amorphous regions, thereby enhancing flexibility, tensile strength, and puncture resistance. These characteristics contribute to the wide application of LLDPE in films, stretch wraps, and pouches (Peacock, 2000).

Plastic pollution has emerged as a worldwide environmental issue due to inadequate recycling and extensive day-to-day usage. Presently, both chemical treatments and bioremediation are employed to address plastic contamination (Omokhagbor Adams et al., 2020). However, chemical approaches often pose environmental risks. As a result, advances in microbial biotechnology have prompted research into isolating microorganisms that degrade plastics from polluted environments such as landfills and waste disposal sites. Bioremediation offers an environmentally friendly solution by utilizing organisms such as bacteria, fungi, algae, or insects (Thirumalaivasan et al., 2024; Zaman et al., 2024). These bioremediation agents are used to break down, eliminate, transform, stabilize, or neutralize plastic pollutants in the environment (Omokhagbor Adams et al., 2020). Recent studies have focused on isolating bacteria capable of metabolizing polyethylene when provided as the sole carbon source (Alamer et al., 2023). Plastic-degrading bacteria constitute a taxonomically diverse group capable of converting synthetic polymers into less harmful compounds through enzymatic reactions. They secrete extracellular and intracellular enzymes such as depolymerases, hydrolases, oxidoreductases, cutinases, amidases, and esterases, which catalyze the hydrolysis or oxidation of chemical bonds within polymer chains (Atanasova et al., 2021; Bahl et al., 2021).

The biodegradation process typically follows several distinct stages. Three major sequential steps can be recognized in plastic degradation: oxidation, depolymerization, and assimilation/mineralization. Initially, biodeterioration occurs as microbes and abiotic factors modify the plastic’s surface, often facilitated by biofilm formation. Oxidation (which may be initiated by abiotic factors such as UV, heat, or mechanical stress, and can also be catalyzed by microbial enzymes) introduces oxygen-containing functional groups (*e.g.*, carbonyls, hydroxyls) into the polymer backbone, increasing hydrophilicity and creating cleavage sites for enzymes (Bahl et al., 2021). At this stage, formation of biofilms likely enhances microbial adhesion to surfaces, thereby increasing the efficiency of enzymatic attack on the polymer. Following oxidation, depolymerization occurs when extracellular enzymes (depolymerases, hydrolases, oxidoreductases) cleave the oxidized polymer into oligomers and monomers that are small enough for transport across the cell envelope. Subsequently, assimilation allows these molecules to traverse the microbial membrane, where they are metabolized into energy and biomass. Finally, mineralization converts assimilated compounds into inorganic end products such as CO and H O (or CH under anaerobic conditions), completing the degradation process (Bahl et al., 2021; Cai et al., 2023).

Plastic-degrading bacteria identified to date differ in their capabilities, with different species and strains exhibiting varying efficiencies in degrading specific polymers. For example, previous studies report that *Bacillus* species such as *B. amylolyticus* and *B. subtilis* reduce the weight of PE samples significantly (Galgali et al., 2002; Nowak et al., 2011). Likewise, *Pseudomonas putida* demonstrated a high PE degradation rate, reducing plastic mass within short incubation periods (Rajandas et al., 2012). A major advancement was the discovery of an *Ideonella sakaiensis* isolate, which produces PETase and MHETase enzymes capable of breaking down PET into assimilable monomers (Yoshida et al., 2016a). Other genera, such as *Rhodococcus*, *Klebsiella*, and *Acinetobacter,* have also been implicated in polyethylene degradation, though often with lower efficiency than *Pseudomona*s and *Bacillus* species (Ghatge et al., 2020; Skariyachan et al., 2016). Even gut bacteria within the insect *Zophobas atratus* have been reported to utilize plastics (LDPE, LLDPE, and EPS) as carbon sources (Zaman et al., 2024).

While many studies have reported diverse bacterial species in plastic degradation, this study is the first to provide a whole-genome analysis coupled with transcriptomic analysis of *Brucella intermedia* strains exhibiting LLDPE-degrading capacity. Unlike commonly reported plastic degraders such as *Pseudomonas* or *Bacillus*, *Brucella* is traditionally recognized as a pathogenic genus (Qureshi et al., 2023). Our findings, therefore, highlight an unexpected ecological role of *B. intermedia*, suggesting that its metabolic repertoire may extend beyond host interactions to include polymer degradation. This not only expands the known diversity of plastic-degrading bacteria but also opens new avenues for exploring unconventional microbial candidates in biotechnological applications.

## 2. Materials and methods

### 2.1 Soil bacteria isolation

Soil samples were collected from the Matuail Sanitary Landfill area in Dhaka, Bangladesh (23°42.97′–23°43.35′ N, 90°26.83′–90°27.2′ E), for isolation of potential LLDPE-degrading bacterial strains ***(SI Figure 1)***. Each sample (1 g) was suspended in 10 mL of sterile 0.9% NaCl solution and subjected to serial dilution. From each dilution, 100 µL was plated onto Minimal Salt Agar (MSA), a carbon-limited selective medium. The composition of MSA was as follows: KH PO (3 g/L), K HPO (0.1 g/L), NaCl (5 g/L), NH Cl (2 g/L), MgSO ·7H O (0.16 g/L), CaCl ·2H O (0.1 g/L), and Agar (2%) with the pH adjusted to 6.5. The medium was sterilized by autoclaving at 121°C for 15 minutes under 15 psi pressure. This medium lacked any organic carbon source. To enable the selection of LLDPE-degrading bacteria, sterilized LLDPE strips were placed on the agar surface after cooling. To sterilize the plastic strips, manufacturer-labeled LLDPE films were washed, cut into uniform strips, and sterilized by soaking in 70% ethanol, as per previous studies (Gupta and Devi, 2020). Controls included: (i) MSA with LLDPE but no bacterial inoculum, and (ii) MSA with inoculum but no LLDPE. Plates were incubated at 37°C for 7 days.

Colonies that emerged at or near the LLDPE film edges were presumed candidate isolates associated with LLDPE-dependent growth. These isolates were individually streaked onto Nutrient Agar (NA) to obtain pure cultures and further cultured in Minimal Salt Broth (MSB), where LLDPE served as the principal carbon source. In each 250 mL Erlenmeyer flask, sterile LLDPE strips were introduced along with 100 mL of MSB and a single bacterial isolate. Controls included MSB with LLDPE only (no inoculum) and MSB with the isolate only (no LLDPE). All systems were incubated at 37°C and 180 rpm for 60 days.

Throughout the incubation, growth was monitored at 7-day intervals using OD readings (data not shown). At the end of the 60-day incubation, these cultures were sub-cultured on NA, and LLDPE films were recovered for Fourier Transform Infrared Spectroscopy (FTIR) and Scanning Electron Microscopy (SEM) to evaluate chemical modification and surface morphological changes. Isolates that showed growth under LLDPE-containing conditions were selected for further biofilm assays, 16S rRNA gene sequencing, whole genome sequencing (WGS), and gene expression analysis.

### 2.2 Fourier Transform Infrared (FTIR) spectroscopy and surface wettability assay

FTIR spectroscopy (IRPrestige-21, SHIMADZU, Japan) was performed on LLDPE films incubated with the isolates for 60 days. Fresh untreated LLDPE was used as the control. FTIR was performed as previously described (Singh et al., 2012).

A hydrophilicity-based droplet spreading assay was also performed on LLDPE films. Sterilized square LLDPE films were incubated with the isolates in MSB for 10 days, with uninoculated MSB-incubated LLDPE used as the control. After incubation, the films were placed on sterile glass Petri dishes and gently blotted with Kimtech Science® Kimwipes™ Delicate Task Wipes (Kimberly-Clark Professional) to remove residual medium. Three 100 µL droplets of distilled water were placed on each film, and images were captured for droplet area measurement. This stage represented the initial surface condition (before surface removal).

The LLDPE surface was then gently wiped using Kimwipes with light pressure to remove the uppermost layer. The droplet spreading assay was repeated following each wiping step. This procedure was performed for four sequential surface-removal stages, and at each stage, the droplet area was measured for both treated and control films using three droplets per sample. Droplet area was quantified from images using ImageJ (Schneider et al., 2012).

### 2.3 Scanning Electron Microscopy (SEM)

SEM was performed on LLDPE films recovered after incubation with isolates X and Y to assess surface morphological changes associated with bacterial exposure. Imaging was carried out using an ultra-high-resolution Schottky Field Emission Scanning Electron Microscope (JSM-7610F, JEOL, Japan). Images were taken after LLDPE samples were incubated in MSB with isolated bacteria for two months. Untreated LLDPE was used as a control. After incubation, LLDPE films were recovered and prepared for SEM as previously described (Taghavi et al., 2021). Images were captured at 3000× magnification; scale bars are provided in the figures.

### 2.4 Biofilm analysis

To assess the biofilm-forming potential of the isolates under both nutrient-rich and hydrocarbon-substituted conditions, qualitative Congo red–based assays were performed following the method of Freeman et al., with modifications (Freeman et al., 1989).

In the first condition (standard Congo red agar, CRA), the medium consisted of Brain Heart Infusion (BHI) broth (37 g/L), sucrose (50 g/L), agar (15 g/L), and Congo red dye (0.8 g/L). For the second condition (modified CRA), hexadecane (100 µL/100 mL) was included as the hydrocarbon-associated substrate, while the concentrations of sucrose, agar, and Congo red dye were kept constant. The corresponding liquid formulation is referred to here as modified Congo red broth (modified CRB). Because hexadecane is poorly miscible with aqueous medium, its distribution was expected to be heterogeneous on solid medium and more available in the liquid phase. Both solid and liquid forms of these media were prepared to observe biofilm formation in agar plates and broth conditions, respectively. This design allowed comparison of biofilm-associated growth under agar- and broth-based hydrocarbon-substituted conditions.

All assays were conducted with appropriate controls. For the hexadecane-supplemented Congo red systems, cultures were monitored at 24 and 48 h to assess time-dependent development of the biofilm-associated phenotype. The interpretation criteria were as follows: colonies exhibiting black coloration or black precipitation on CRA plates or in broth were considered positive for biofilm-associated growth, whereas colonies remaining red or pink without black precipitation were classified as negative under the tested conditions.

In addition to qualitative Congo Red assays, biofilm formation was further examined using direct observation of LLDPE-associated growth and SEM to evaluate bacterial attachment and biofilm architecture associated with LLDPE. For SEM analysis, two experimental systems were established for each isolate. In the first system, modified CRA was prepared as described previously, except that sterilized LLDPE films were used as the hydrocarbon substrate instead of hexadecane to allow direct observation of biofilm formation on the plastic surface. In the second system, isolates were cultured in modified CRB containing LLDPE to facilitate imaging of bacterial aggregate formation in the liquid phase after exposure to plastic. All systems were incubated at 37°C under identical conditions. Following incubation, LLDPE films from the first system were carefully recovered, gently washed with sterile phosphate-buffered saline (PBS) to remove loosely attached cells, and processed according to the previously described SEM preparation protocol. For the second system, biofilm biomass formed in the broth was collected and prepared for SEM imaging. For SEM-based analysis of biofilm-associated growth, LLDPE films and recovered broth biomass were processed and imaged using the same protocol described in Section 2.3. This approach enabled comparative evaluation of biofilm formation and structural characteristics under plastic-associated, nutrient-limited conditions. In addition, LLDPE films placed on modified CRA were monitored over time to visualize progressive surface colonization under plastic-associated conditions.

### 2.5 Whole genome sequencing and assembly

Raw Oxford Nanopore Technologies (ONT) reads were processed using a custom Snakemake workflow. Initial taxonomic classification was performed using Kraken2 (v2.0.8-beta), and reads assigned to predefined contaminant taxa were removed using seqkit grep (SeqKit, v2.10.0) (Shen et al., 2024; Wood et al., 2019). Read quality was evaluated using NanoPlot (v1.44.1), and filtering was conducted with NanoFilt (v2.8.0), retaining reads with minimum length of 1000 bp and average quality ≥10 (De Coster et al., 2018). Filtered reads were assembled de novo using Flye (v2.9.5) with an estimated genome size of 5 Mb (Kolmogorov et al., 2019). Genome/chromosome circularity status was inferred from Flye output. Assembly polishing was performed in two stages: three iterative rounds of Racon (v1.5.0), followed by consensus refinement with Medaka using the model r1041_e82_260bps_hac_v4.1.0. Assembly completeness was evaluated with BUSCO (v5.8.3) 5 using the bacteria_odb12 dataset (Manni et al., 2021). Assembly quality and structural metrics were further assessed using QUAST (v5.0.2). Sequencing depth was estimated by aligning filtered reads to the polished assembly using Minimap2 (v2.26), followed by sorting, indexing, and coverage computation with SAMtools (v1.21) using samtools coverage.

### 2.6 Taxonomic Assignment and Reference Genome Retrieval

Whole-genome assemblies of the two bacterial isolates, designated X and Y, were initially taxonomically classified using Kraken2 v2.0.8-beta with the pluspf16 Kraken2 database. Both genomes were confidently assigned to the genus *Brucella*. Independently, 16S RNA sequences of both bacterial isolates also indicated that both isolates belonged to the *Brucella* genus. To identify appropriate reference genomes for comparative analysis, all complete genome assemblies of the genus *Brucella* were retrieved from the NCBI RefSeq database (date: May 14, 2025). Only assemblies at the “Complete Genome” level were included to ensure consistency and high sequence quality (n = 231).

### 2.7 Average Nucleotide Identity (ANI) Screening & Isolate Difference Assessment

Species-level classification was refined using FastANI v1.1, which calculates pairwise Average Nucleotide Identity (ANI) between query and reference genomes (Jain et al., 2018). Genomes of isolates X and Y were also compared with each other using reciprocal Minimap2–Medaka (v2.0.1, model: r1041_e82_260bps_hac_variant_v4.1.0_model_pt.tar.gz) variant calling and whole-genome alignment with MUMmer v3.23 (dnadiff/nucmer).

### 2.8 Whole genome comparison

Whole-genome relationships among representative *Brucella* isolates were assessed using Mashtree (v1.4.6) (Katz et al., 2019). Complete and representative *Brucella* genomes from NCBI, spanning both pathogenic and environmental lineages, were analyzed alongside the study isolates X and Y *(SI Table 1)*. The resulting distance matrix was converted to a neighbor-joining tree, midpoint-rooted, and visualized in ETE3 (v3.1.3) (Huerta-Cepas et al., 2016), with branch-lengths square-root–scaled and species names mapped from NCBI metadata.

For focused genome comparison, each landfill isolate (X and Y) was mapped to the closest high-quality *B. intermedia* reference genome (GCF_048398275.1; hospital-wastewater origin). For every annotated CDS in the reference, their presence-absence in the landfill isolates and relocation status were checked. A CDS was called absent if <80% of its length was covered at the native locus and no relocated match was detected in the isolate assembly (≥80% query coverage, ≥90% amino acid identity). Consecutive absent CDS located <5 kb apart on the same reference contig were merged into genomic “accessory islands.” Genomic span, GC content, number of CDS, and representative annotations were checked for each of the islands. These islands were visualized as depth dropouts in a circular genome plot, rendered from whole-genome alignments and coverage tracks using minimap2, samtools, and pyCirclize. Complete calling criteria, coordinates, and functional summaries are provided in *SI Note 1* and *SI Table 2*.

### 2.9 Pairwise variant identification and visualization

Single nucleotide polymorphisms (SNPs) and small insertions/deletions (indels) between the landfill isolates (X and Y) were identified using Snippy v4.6.0 (Seemann, 2025a), with the Y isolate assembly as the primary reference. Variants were parsed from the resulting VCF file, classified as SNP or indel, binned by a 100 bp genomic coordinate, and plotted as a stacked lollipop plot using Matplotlib v3.9 using a custom *in-house* Python script. In the stacked lollipop plot, each stick represents local variant pile-up along the two chromosomes of the reference genome(s) (GCF_043898275.1 for *Figure 3C* and the Y isolate genome for *SI Figure 5*). SNPs and indels were visualized together as stacked lollipop plots to illustrate overall genome-wide variation. In *SI Figure 5*, these were shown separately to distinguish variant types across the two chromosomes. The region absent in X (2.255–2.275 Mb) was further examined using the corresponding locus from the closest published *B. intermedia* reference genome (GCF_043898275.1; *SI Figure 5C*); gene models were extracted directly from its NCBI GFF annotation file to indicate gene content and putative functions.

### 2.10 Identification of plastic-degrading genes

Genome assemblies (FASTA format) were screened with an in-house workflow. Protein-coding genes were predicted using Prokka v1.14.6 (default settings). The resulting proteins were queried against PlasticDB (May 18, 2025 release) with BLASTP v2.16.0 (E-value ≤ 1 × 10^O5^) (Gambarini et al., 2022). A match was accepted when (i) amino-acid identity ≥ 30 %, (ii) query coverage ≥ 60 %, and (iii) subject coverage ≥ 60 %. BLAST hits were parsed to extract enzyme annotations from the target sequence identifiers, which encoded enzyme names in a delimiter-separated format. Enzyme counts were summarized per genome, and individual summaries were aggregated into a master count matrix for comparative analysis.

### 2.11 Resistome and virulence factor prediction and antimicrobial susceptibility testing

Antimicrobial resistance and virulence-associated genes were identified using ABRicate on the Galaxy (U.S.) server (Afgan et al., 2018; Seemann, 2025b) with the ResFinder (Zankari et al., 2012) and VFDB databases (≥70% identity, ≥80% coverage) (Chen et al., 2016). Additional resistance analysis was performed using the Comprehensive Antibiotic Resistance Database (CARD) via the Resistance Gene Identifier (RGI) web interface (Alcock et al., 2023) with default settings. Phenotypic antimicrobial susceptibility testing was performed using the VITEK^®^ 2 system (bioMérieux, Marcy l’Etoile, France) with AST-N405 cards according to the manufacturer’s instructions.

### 2.12 Biofilm-forming gene prediction

A curated dataset of 63 biofilm-associated proteins was compiled from experimentally validated systems across diverse microorganisms, including *Staphylococcus aureus*, *Pseudomonas aeruginosa*, *Escherichia coli*, *Vibrio cholerae*, *Bacillus subtilis*, *Enterococcus faecalis*, and *Candida albicans,* through a literature survey (*SI Table 3*). These reference proteins represent core components of adhesion, matrix biosynthesis, quorum-sensing, and regulatory networks involved in biofilm formation. Protein sequences were obtained from UniProt and used as a BLASTp query set against the predicted proteomes of *Brucella* landfill isolates X and Y.

### 2.13 Transcriptomic data collection and analysis

LLDPE was incubated with isolate X in MSB for 30 days. Harvested bacterial cells were immediately treated with Qiagen RNAprotect Bacteria Reagent (Cat. No. 76506) to stabilize RNA. The stabilized cell pellets were transported to DNA Solution Ltd. (Dhaka, Bangladesh) under refrigerated conditions (4°C) and subsequently stored at −80°C until further processing. Total RNA was extracted using the Qiagen RNeasy Mini Kit (Cat. No. 74104) according to the manufacturer’s instructions. For transcriptome analysis, sequencing libraries were prepared using the Illumina Stranded Total RNA Prep, Ligation with Ribo-Zero Plus kit (Cat. No. 20040525), which enables ribosomal RNA depletion and strand-specific library construction. The prepared libraries were sequenced on the Illumina NextSeq 2000 platform using paired-end chemistry (2 × 150 bp). Paired-end Illumina RNA-seq reads were adapter- and quality-trimmed using fastp (Chen et al., 2018), then aligned to the reference genome with Bowtie2 (Langmead and Salzberg, 2012). Alignments were processed (BAM conversion, sorting, indexing) with SAMtools (Li et al., 2009), and downstream quantification was restricted to high-confidence reference alignments (MAPQ ≥ 20). Gene-level read counts were generated with featureCounts (Subread) (Liao et al., 2014), and expression was reported as TPM (transcripts per million; length- and library-size normalized) and visualized as log_2_(TPM+1) (Wagner et al., 2012).

## 3. Results and discussion

### 3.1 Landfill microbiota harbor LLDPE-degrading bacterial isolates

Following a 7-day incubation on MSA plates containing LLDPE, four to five distinct colony morphotypes were observed growing adjacent to the LLDPE film margins. In contrast, the negative control plate containing LLDPE without soil inoculum exhibited no visible growth, confirming the sterility of the LLDPE and the selectivity of the medium. Plates containing only the soil sample (without LLDPE) also showed some colonies, as expected due to the presence of organic matter in the soil. Therefore, to further validate potential LLDPE degraders, all colonies from the MSA plates with both LLDPE and inoculum were individually subcultured on Nutrient Agar to obtain pure cultures. These pure cultures were subsequently inoculated individually into Minimal Salt Broth (MSB) with LLDPE as the principal carbon source. Growth was monitored every 7 days by OD measurements (data not shown). After 60 days, only two isolates showed sustained growth increases relative to their initial OD values, suggesting active proliferation under LLDPE-containing conditions.

To reduce false positives and assess long-term viability, all MSB cultures were subsequently subcultured onto NA after day 60. In the systems containing only the isolates (no LLDPE), no growth was observed beyond day 7, indicating that the bacteria could not survive in the absence of an external carbon source. Similarly, the uninoculated LLDPE control flasks remained sterile throughout the 60-day period. These results support the conclusion that only two isolates were able to persist and grow under these conditions in the presence of LLDPE. Thus, the initial growth observed during the first incubation may have been driven by residual organic matter in the soil or media or by carryover of non-growing biomass, potentially leading to initial false positives. However, the extended incubation in MSB, a strictly inorganic medium, and the subsequent lack of growth in control systems supported the identification of the two candidate LLDPE-degrading strains, designated isolates “X” and “Y”. The recovered LLDPE films from these two systems were subjected to FTIR and SEM analyses to assess chemical modification and surface morphological changes.

The selective isolation of only two degraders highlights that microorganisms with demonstrable LLDPE-degrading capacity represent only a minority of the cultivable community recovered from landfill-associated samples. Similar low-yield recovery has been reported in previous screening studies (Nademo et al., 2023), where only a subset of soil or waste-site isolates showed measurable polyethylene utilization or degradation under selective conditions. Moreover, extended incubation in MSB medium is particularly important, as it reduces the risk of false positives that may arise from residual organic matter and helps enrich for strains capable of sustained growth under polymer-associated, carbon-limited conditions. Furthermore, the observed persistence and growth under LLDPE-containing, carbon-limited conditions suggest specialized metabolic adaptations. Similar findings have been reported in other species, where oxidative enzymes and biofilm-associated proteins enabled survival under carbon-limited, polymer-dependent conditions (Skariyachan et al., 2016). This suggests that isolates X and Y may employ comparable oxidative and/or hydrophobic interaction strategies. Ecologically, as plastic pollution accumulates in the environment, plastic degrading bacteria may gain increased access to substrates, enabling greater energy acquisition, growth, and persistence. More broadly, landfill environments, rich in hydrocarbons and recalcitrant polymers, can act as hotspots for the emergence of microbial populations with specialized plastic-degrading capabilities.

### 3.2 Activities of plastic-degrading bacterial isolates induce chemical changes in LLDPE

Microbial degradation of LLDPE is typically accompanied by chemical modifications in the polymer backbone (Gautam et al., 2007). To determine whether the landfill isolates induced such changes, FTIR analysis was performed on LLDPE films after 60 days of incubation. The untreated control exhibited the characteristic polyethylene C–H stretching bands, along with a weak band at ∼1020 cm□¹ (C–N stretch), possibly associated with additives or processing residues (*Figure 1*). In contrast, LLDPE films treated with isolates X (*Figure 1*) and Y (*SI Figure 2*) showed the appearance of additional bands assigned to oxygen-containing functional groups that were absent in the control. These included carbonyl stretching bands (1550–1720 cm□¹), C–O stretching bands (1250–1300 cm□¹), and a broad O–H stretching band (3000–3670 cm□¹). The appearance of these functional groups is consistent with oxidative modification of the polymer surface and suggests that bacterial activity altered the chemical structure of LLDPE under carbon-limited conditions, where the polymer was available as the principal carbon source.

**Figure 1.**
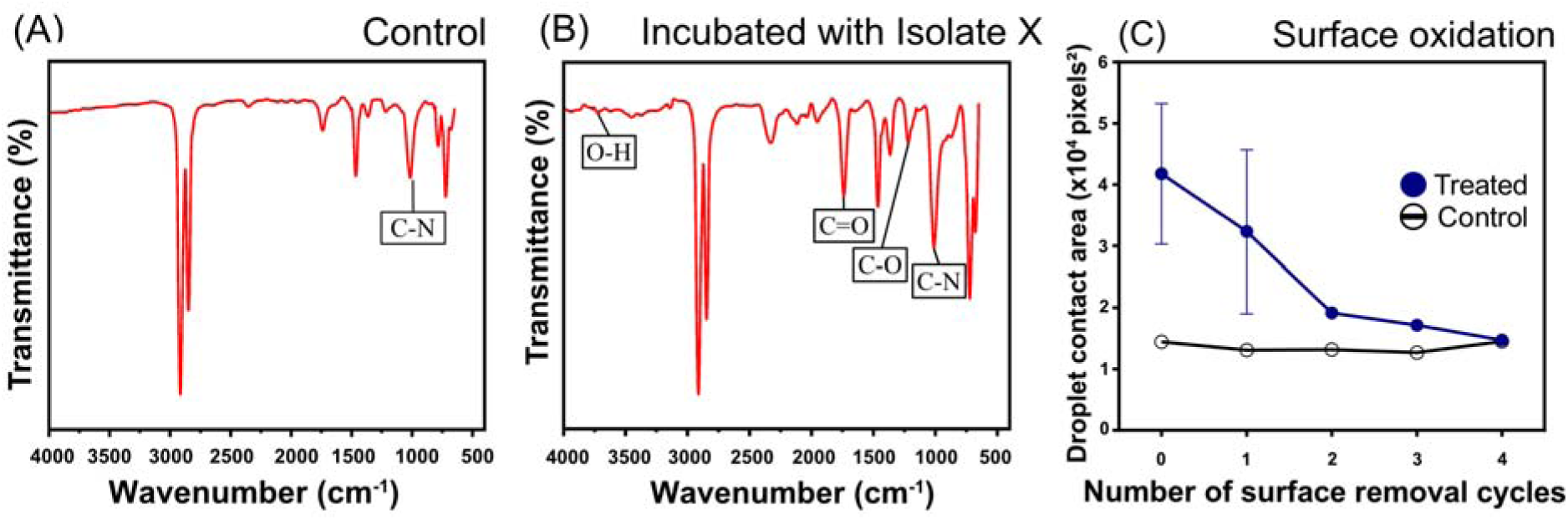
Chemical modification of LLDPE following exposure to landfill *Brucella intermedia* isolate X. **(A)** Fourier Transform Infrared (FTIR) spectrum of untreated control LLDPE, showing the characteristic polymer bands and a weak band near ∼1020 cm□¹, possibly associated with additives or processing residues. **(B)** FTIR spectrum of LLDPE after treatment with isolate X, showing additional bands assigned to O–H (3000–3670 cm□¹), C=O (1550–1720 cm□¹), and C–O (1250–1300 cm□¹) functional groups, consistent with oxidative modification of the polymer surface. **(C)** Distilled water droplet contact area on control and treated LLDPE across successive surface removal cycles. Treated LLDPE showed a larger droplet contact area than the control, indicating increased surface wettability. This difference decreased with successive surface removal cycles, and by the fourth cycle, the treated and control samples showed similar droplet areas. Droplet area was quantified using ImageJ (Schneider et al., 2012).

The observed FTIR peaks correspond to oxygenated intermediates commonly associated with polyethylene oxidation (Khumthong et al., 2025; Socrates, 2004). The O–H stretching band suggests the formation of hydroxylated or alcohol intermediates, while the carbonyl signal near ∼1700 cm□¹ likely represents ketones, aldehydes, or carboxylic acids formed during polymer oxidation. The presence of C–O stretching bands further supports the formation of ester- or ether-like intermediates. Similar oxidative signatures have previously been reported as indicators of microbial polyethylene degradation (Khumthong et al., 2025). The incorporation of these oxygen-containing groups increases the polarity of the otherwise hydrophobic polyethylene backbone, thereby increasing surface wettability and potentially enhancing susceptibility to subsequent cleavage or downstream processing.

To further assess whether these changes were concentrated at the polymer surface, a hydrophilicity-based droplet spreading assay was conducted (*SI Figure 3*). Water droplets placed on bacteria-treated LLDPE films exhibited significantly larger spreading areas than those on untreated controls, indicating increased surface wettability following incubation, consistent with oxidation-associated introduction of polar groups onto the polyethylene surface (Holmes-Farley et al., 1985) (*Figure 1, SI Figure 3*). When the outermost polymer layer was gradually removed through sequential surface wiping, the droplet spreading area progressively decreased (*Figure 1*). After four removal stages, the droplet behavior resembled that of untreated control films (*Figure 1, SI Figure 3*), suggesting that the wettability change was primarily localized to the bacteria-exposed outer surface layer. This pattern is consistent with surface-associated oxidative modification, although contributions from biofilm-associated material at the exposed surface remain possible (see Section 3.6). Taken together, these data support the interpretation that bacterial exposure introduced polar surface-associated modifications to the LLDPE, thereby contributing to increased wettability.

The emergence of O–H and C=O bands in the treated samples is consistent with the oxidation–depolymerization–assimilation model of plastic biodegradation (Atanasova et al., 2021; Khumthong et al., 2025). In this mechanism, oxidative enzymes such as oxidases or peroxidases introduce oxygen-containing groups into the polymer backbone, which subsequently facilitates chain scission and conversion of the polymer into smaller oligomers that can be metabolized by microorganisms (Atanasova et al., 2021). Although both isolates exhibited similar oxidative signatures, qualitative differences in band intensity and droplet spreading patterns may reflect variation in the extent of surface modification or in the underlying enzymatic response between the strains. Collectively, these findings demonstrate that isolates X and Y induce chemical modification of LLDPE consistent with surface oxidation, a crucial initial step in microbial polyethylene degradation.

### 3.3 Activities of plastic-degrading bacterial isolates alter LLDPE surface morphology

Because chemical changes occur at the molecular scale, we next asked whether isolate-mediated chemical modification of LLDPE was accompanied by detectable changes in surface morphology. To address this, we directly visualized untreated and bacteria-exposed LLDPE films using SEM. Scanning electron micrographs revealed clear differences in the surface morphology of LLDPE films incubated with isolates X (*Figure 2*) and Y (*SI Figure 4*) compared to the untreated control. After 60 days of incubation, the control film maintained a mostly smooth surface, whereas the treated samples displayed roughened and irregular surface morphology, including localized fissures and fragmentation-like features. Such morphological alterations are consistent with previous observations on microbial plastic degradation, where bacterial colonization and enzymatic action gradually modify and erode the polymer surface (Kyaw et al., 2012; Shah et al., 2008). The roughness and cracking observed on the LLDPE surfaces likely result from bacterial exposure–associated oxidative modification of the polymer backbone, as supported by our FTIR spectroscopy findings (*Figure 1*). These changes may, in turn, promote further microbial attachment by increasing surface irregularity and exposing chemically modified regions of the polymer, thereby potentially facilitating subsequent depolymerization steps. Similar SEM-based observations have been reported in other bacterial strains grown on polyethylene films, where extensive pitting, grooves, and cavities were interpreted as signatures of microbial surface attack (Skariyachan et al., 2016).

**Figure 2.**
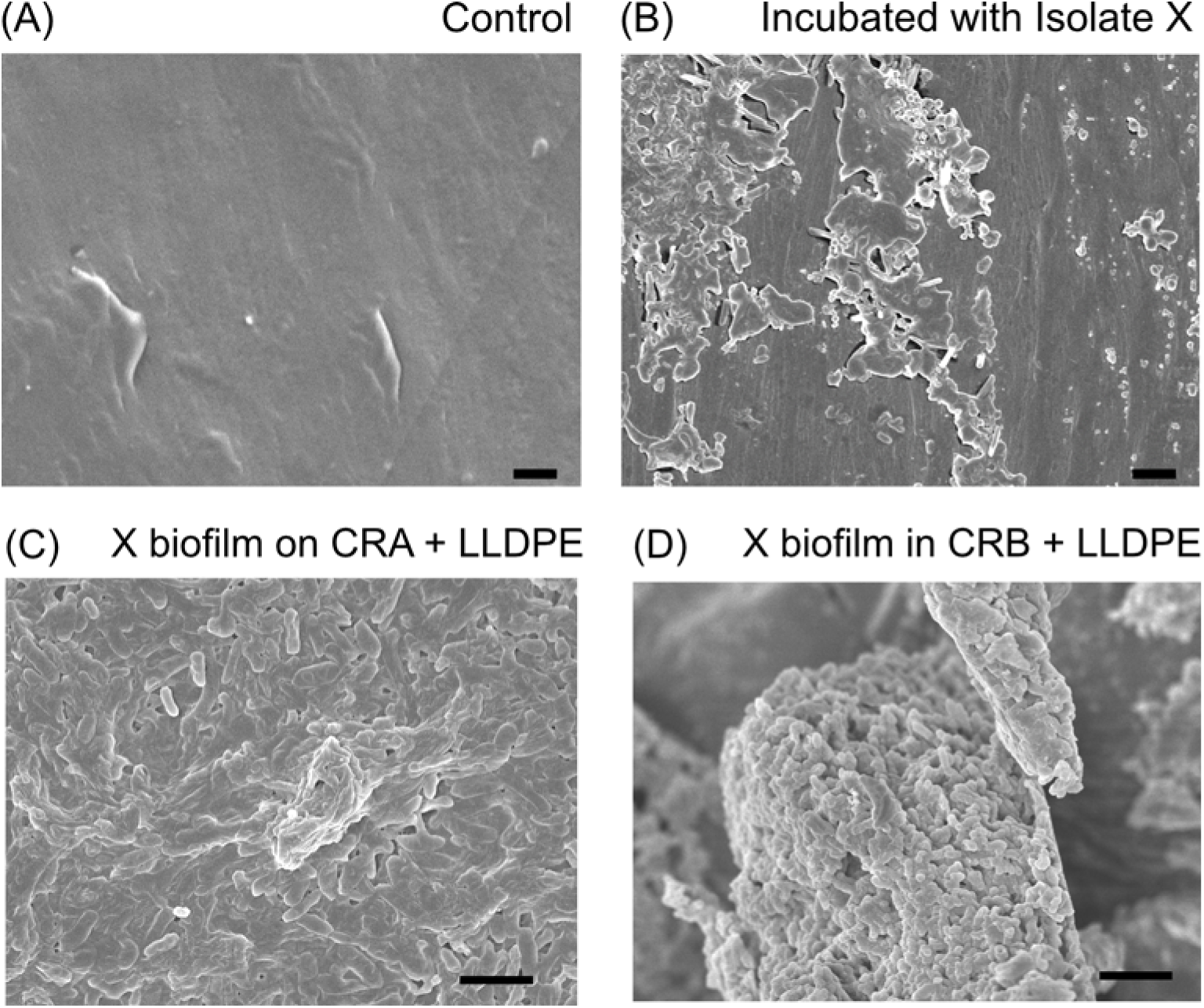
Scanning electron micrograph analysis of LLDPE surface alterations and bacterial responses associated with LLDPE exposure using isolate X. **(A)** Untreated control LLDPE showing a mostly smooth surface. **(B)** Cleaned LLDPE recovered after bacterial incubation, showing a roughened and irregular surface compared with the untreated control. **(C)** Dense biofilms are embedded within an extracellular polymeric substance (EPS), covering large areas of the treated plastic surface. **(D)** Dense bacterial aggregates recovered by centrifugation from broth cultures after incubation with LLDPE, showing compact clump-like biomass with matrix-like morphology (for raw data, see *SI* Figure 3). Scale bars represent 3 µm.

### 3.4 Genomic characterization and LLDPE-responsive transcriptional profiling in the landfill isolates

For species-level identification and a clear understanding of the genetic basis of LLDPE-degrading capacity, we performed whole-genome shotgun sequencing of the two landfill isolates. DNA from pure colonies of each isolate was sequenced, and the reads were assembled and annotated for detailed genomic characterization. Taxonomic classification using Kraken2 indicated that both bacterial isolates belong to the *Brucella* genus. For species identification, we compared assembled X and Y isolate genomes against all available high-quality *Brucella* reference genomes. Both X and Y isolates showed high average nucleotide identity (ANI) scores against previously reported *B. intermedia* genomes (*SI Table 6*). To further delineate isolate taxonomic identities, we performed whole-genome phylogenetic analysis with representative *Brucella* genomes. Results of the phylogenetic analysis showed that both X and Y isolate genomes cluster with other previously sequenced *B. intermedia* isolates (*Figure 3A*). Interestingly, the environmental isolates included in our analysis clustered together (*Figure 3A; SI Table 6*), confirming their close genomic affinity and environmental origin. The whole-genome phylogenetic analysis reveals a clear separation between classical pathogenic and environmental clades, reflecting the evolutionary divergence between host-adapted and free-living *Brucella* lineages (*Figure 3A*).

**Figure 3.**
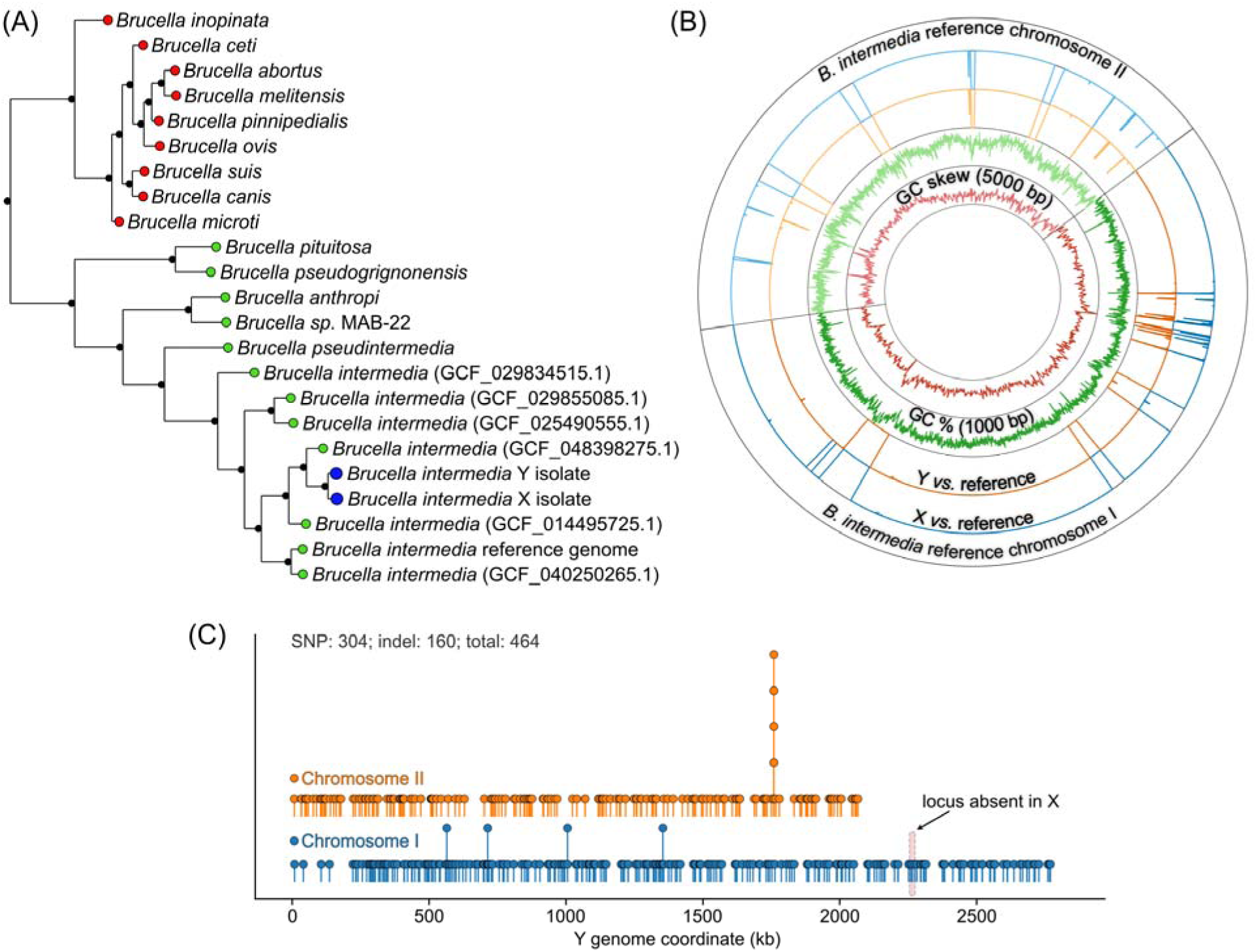
Genomic and phylogenomic analysis of landfill-derived *Brucella intermedia* isolates X and Y. (**A**) *Isolates X and Y cluster with environmental Brucella intermedia clades.* Whole-genome phylogeny of the *Brucella* genus. Environmental species (green-filled circle at the tip) and pathogenic species (red-filled circle at the tip) form distinct clades; isolates X and Y (blue-filled large circle at the tip) cluster with *B. intermedia*. Tree visualized with a custom root for clarity and square-root-scaled branch lengths. (**B**) *Landfill isolates are distinct from their nearest sequenced B. intermedia neighbor.* Circos plot: the *B. intermedia* reference genome (GCF_048398275.1; outer ring), X vs. reference and Y vs. reference comparison tracks (middle rings), and GC% and GC skew tracks (inner rings). Within the X and Y comparison tracks, dark and light shades denote chromosomes I and II, respectively. White gaps in the X/Y rings mark “accessory genomic islands” that are present in the hospital-wastewater reference but missing from one or both landfill isolates. (**C**) *X and Y isolates are genomically distinct.* Genome-wide single nucleotide polymorphisms (SNPs) and small indels between isolates X and Y (visualized using isolate Y genome as the reference coordinate system). Variants were visualized as stacked lollipop plots across the two chromosomes, where vertical offset markers represent overlapping or closely spaced variants (pile-ups). A ∼19.8 kb region, exclusive to isolate Y and absent in isolate X, is delineated by the red-shaded box.

The whole genome phylogenetic tree showed that both X and Y isolates are close to an environmental *B. intermedia* isolate (GCF_048398275.1; a hospital wastewater isolate from Ghana), with ANI scores of 99.56 and 99.59, respectively (*SI Table 6*). Such a high ANI is typically observed among isolates of the same species or very closely related strains. To directly visualize similarities/differences between the GCF_048398275.1 isolate and X-Y isolates, we compared identities of the landfill isolates *vs.* the nearest reference genome. Whole genome alignment showed that X and Y isolates are largely syntenic but lack discrete multi-gene cassettes that appear in the reference wastewater isolate (*Figure 3B*). These “accessory islands” include phage/plasmid modules and a high-GC (∼0.62–0.63) stress/repair block (∼2.16–2.22 Mb in the reference) encoding MarR/LysR-family regulators and MsrB, all absent from both landfill isolates (*SI Note 1*).

To characterize the difference between the two landfill isolates, we checked the ANI scores when X and Y are compared with each other. Our analysis revealed that X and Y share a high degree of similarity with ANI scores >99.98. To check whether X and Y are clonally identical, isolate X was directly aligned with isolate Y (*Figure 3C*). A total of 304 single-nucleotide polymorphisms (SNPs) and 160 indels were identified. These variations are uniformly distributed across the two *B. intermedia* chromosomes, without evidence of localized hypermutation. Notably, we also detect a smaller locus present in Y but missing in X that encodes a predicted two-component histidine kinase/sigma factor pair and envelope glycosylation genes (*Figure 3B, SI Figure 5C, SI Table 2*). Together, these differences confirm that the two landfill isolates are closely related but genomically distinct lineages of *B. intermedia*, consistent with microevolution within a shared environmental niche.

To understand the molecular basis of plastic-degrading capacity, we compared these isolates’ annotation-derived proteomes against a manually curated set of plastic-degrading proteins (Gambarini et al., 2022). For both isolates, 42 putative plastic degraders were identified (*SI Table 4*). These enzymes represented all key steps in plastic biodegradation (namely, oxidative activation, depolymerization, and assimilation/mineralization). To check whether these genes are significantly enriched vis-à-vis the *Brucella* reference genome background, we determined baseline counts (mean ± standard deviation) for each of these enzymes within publicly available *Brucella* genomes and compared them with identified plastic-degrading genes from X and Y isolates (*Table 1*). Relative to the reference *Brucella* genomes, isolates X and Y were strongly enriched in oxidative activation capacity, moderately enriched in depolymerization enzymes, and showed a near-baseline assimilation potential. Our observations mirror previous reports on plastic-degrading enzyme category expansions in known plastic degraders [*Rhodococcus* spp. (oxidative activation) and *Ideonella* (depolymerization)] (Santo et al., 2013; Yoshida et al., 2016b; Zampolli et al., 2023).

**Table 1:** Distribution of plastic-degradation–associated enzyme categories in reference *Brucella* genomes (n = 231) and the landfill isolates. Reference genome means and standard deviations (SD) were calculated across 231 genomes. Isolate values represent the number of detected homologs per genome; isolates X and Y showed identical counts. Z-scores are (count−mean)/SD; NA indicates SD = 0.

| Enzyme Category | Enzyme | Reference *Brucella* genomes (n=231) |  | *Brucella* landfill isolates |  |
| --- | --- | --- | --- | --- | --- |
|  |  | Mean | Standard Deviation (SD) | Count | Z-score |
| Oxidative activation | Copper oxidase | 0.01 | 0.11 | 1 | 8.7 |
|  | Laccase | 0.03 | 0.16 | 0 | -0.16 |
| Depolymerization | PHA depolymerase | 0.04 | 0.19 | 0 | -0.2 |
|  | PLA depolymerase | 1 | 0 | 1 | NA |
|  | PU esterase | 0 | 0.07 | 0 | -0.07 |
|  | Polyamidase | 0.1 | 0.37 | 1 | 2.46 |
|  | Polyesterase | 1 | 0 | 1 | NA |
| Assimilation-mineralization | 3HV dehydrogenase | 13.45 | 3.77 | 20 | 1.74 |
|  | Carboxylesterase | 0.03 | 0.16 | 0 | -0.16 |
|  | Hydrolase | 0.62 | 0.49 | 0 | -1.28 |
|  | Lipase | 0.21 | 0.75 | 0 | -0.28 |
|  | Nylon Oligomer Degrading Enzyme | 1.25 | 0.5 | 1 | -0.5 |
|  | Nylon hydrolase | 0.03 | 0.16 | 0 | -0.16 |
|  | PEG aldehyde dehydrogenase | 13.01 | 0.89 | 13 | -0.01 |
|  | PEG dehydrogenase | 4.69 | 0.66 | 4 | -1.04 |

Genomic profiling of the 42 candidates further clarified the stage-wise distribution of putative functions (*Table 1*). Among the oxidative activation genes, copper oxidase was strongly overrepresented in both landfill isolates (Z = 8.7), suggesting a prominent copper-dependent oxidative toolkit to initiate polymer attack. Copper oxidases can catalyze electron transfer reactions that introduce carbonyl and hydroxyl functionalities on hydrocarbon chains, increasing hydrophilicity and facilitating subsequent cleavage (Santo et al., 2013). This enrichment is consistent with our FTIR observations that prolonged incubation of LLDPE with the landfill isolates yields strong carbonyl and hydroxyl peaks (*Figure 1*), hallmarks of oxidative modification. Depolymerization genes were moderately enriched, including PLA depolymerase, polyesterase, and polyamidase with counts above or near reference means (*Table 1*), supporting the potential for hydrolytic cleavage of oxidized polymer fragments and/or related aliphatic substrates (Guo et al., 2024; Shalem et al., 2024). In contrast, assimilation/mineralization categories were near baseline by gene inventory, which is expected for broadly conserved central metabolic and oxidoreductase functions across *Brucella* genomes.

Importantly, gene inventory enrichment reflects genomic potential, whereas transcript abundance under LLDPE growth reflects condition-specific pathway engagement. To assess whether the predicted plastic-degradation candidates are transcriptionally active under LLDPE exposure, we performed RNA-seq on isolate X grown under the LLDPE condition (three biological replicates) and quantified expression as TPM (transcripts per million). Within the 42-gene candidate panel, stage-structured expression showed a strong signal in oxidative-attack and assimilation-associated candidates (*Figure 4A*), consistent with a model in which oxidative activation is accompanied by transcriptional engagement of downstream processing and metabolite-utilization functions during LLDPE growth. Candidate-gene expression profiles displayed replicate-to-replicate heterogeneity, with LLDPE-1 and LLDPE-2 more similar to each other than to LLDPE-3 (*Figure 4B*), and with corresponding shifts in the relative allocation of candidate-panel TPM across functional stages (*Figure 4C*). At the enzyme-family level, a small set of families accounted for the highest summed candidate expression under LLDPE growth, including copper oxidase (oxidative attack) and PEG aldehyde dehydrogenase and 3HV dehydrogenase homologs (assimilation) (*Figure 4D*). Depolymerization candidates were present and expressed but represented a smaller fraction of the candidate-panel transcript pool than oxidative and assimilation functions in these libraries (*Figure 4C*), which may reflect temporal staging, differing regulatory control, or lower transcript requirements for hydrolytic steps at the sampled timepoint. Although transcript abundance does not directly measure enzyme activity, these results provide independent support that multiple predicted plastic-degradation–associated functions are expressed under LLDPE growth, complementing the genome-derived enrichment patterns (*Table 1; SI Table 4*) and aligning with chemical evidence of oxidative modification (*Figure 1*).

**Figure 4.**
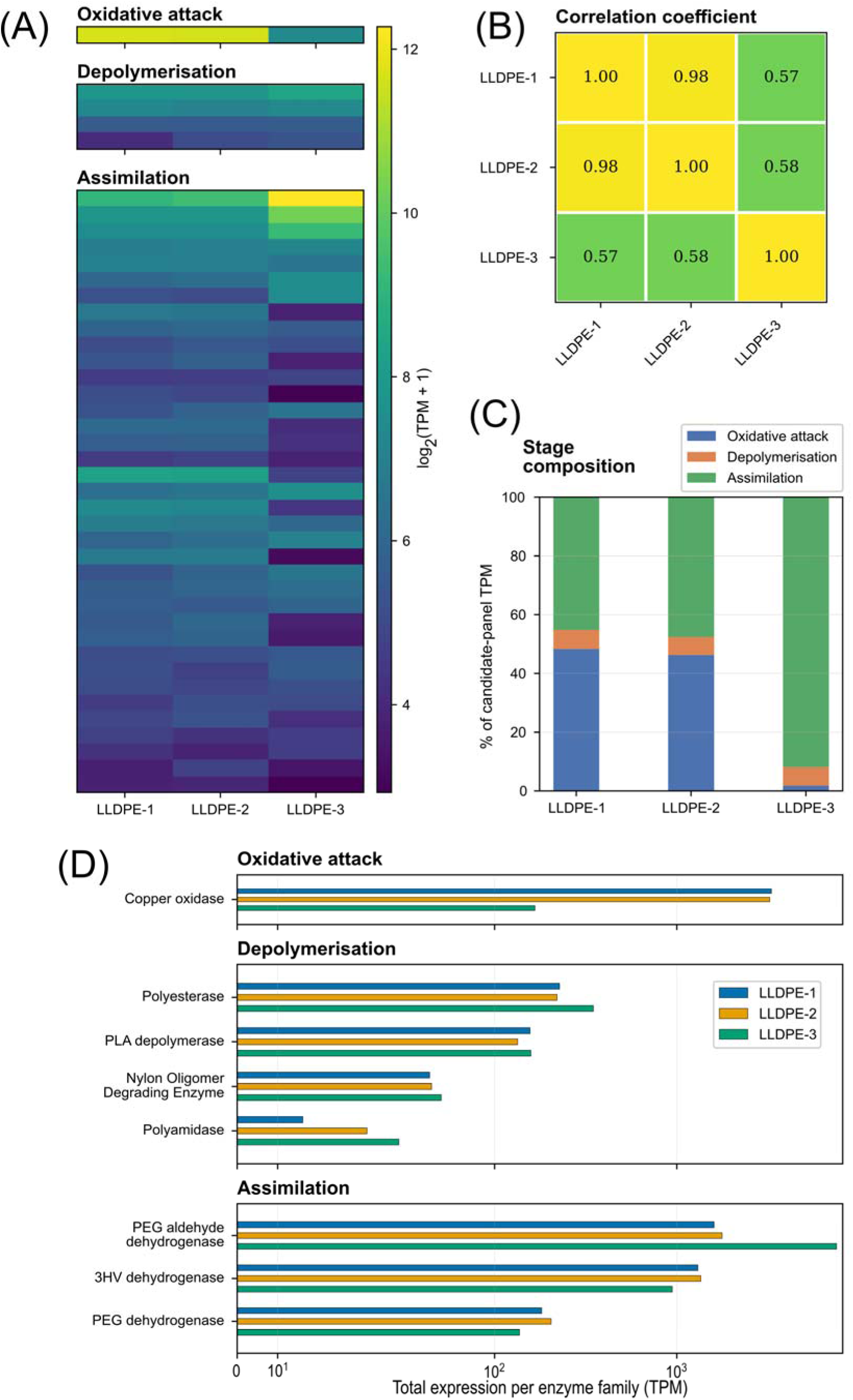
Stage-resolved expression patterns of candidate LLDPE degradation and assimilation genes in landfill Brucella intermedia isolate X. (**A**) *Oxidative attack and assimilation candidates are highly expressed under LLDPE growth, consistent with putative roles in polymer attack and metabolite utilization in B. intermedia (isolate X).* Heatmap of 42 candidate plastic degrading genes grouped by functional stage (oxidative attack, depolymerisation, assimilation). Expression is quantified as TPM (transcripts per million; length-and library-size normalized) and shown as log_2_(TPM+1) across three LLDPE-grown transcriptomes (LLDPE-1 to LLDPE-3). The heatmap highlights strong expression of candidate genes assigned to oxidative attack and assimilation categories under LLDPE growth. (**B**) *Candidate-gene expression profiles are more similar between LLDPE-1 and LLDPE-2 than to LLDPE-3.* Pearson correlation coefficients (*r*) computed on log_2_(TPM+1) values across the 42-candidate gene panel; cell values report *r*. (**C**) *Relative stage composition differs across replicates, with oxidation/assimilation contributions varying across LLDPE-1 to LLDPE-3.* Stacked bars show the percentage contribution of oxidative attack, depolymerisation and assimilation to the total TPM summed across the 42 candidate genes within each replicate. (**D**) *A small set of enzyme families accounts for the highest candidate-panel expression under LLDPE growth.* At the enzyme-family level, copper oxidase (oxidative attack) and PEG aldehyde dehydrogenase and 3HV dehydrogenase homologs (assimilation) show the highest summed expression among the candidates. For each stage, TPM values were summed across homologous candidate genes assigned to the same enzyme family and plotted on a symmetric log scale to accommodate the dynamic range in abundance.

Together, these genomic and transcriptomic analyses support the classification of isolates X and Y as closely related environmental *B. intermedia* lineages with distinct but highly similar genomes and with a conserved repertoire of candidate functions plausibly involved in LLDPE oxidation, depolymerization, and downstream assimilation.

### 3.5 Resistome and virulence-related features of the landfill isolates

Previous studies have reported plastic-degrading capacities of multiple *Brucella* spp., including *B. intermedia* (Alamer et al., 2023; Srivastava et al., 2025), which aligns with the combined genomic and transcript-level evidence presented here and suggests that these strains may be adapted to environmental niches rich in hydrocarbon or polymer substrates. Because environmental deployment or handling of plastic-degrading bacteria also raises biosafety considerations, we next examined the antimicrobial resistance and virulence-associated features of the two landfill isolates.

Combined resistome profiling using the Comprehensive Antibiotic Resistance Database (CARD; RGI; Alcock et al., 2023) and ResFinder (ABRicate; Seemann, 2020) revealed that both X and Y genomes possess an almost identical antimicrobial resistance repertoire. Each genome encodes three RND-type efflux pumps (adeF-like), a small multidrug resistance (SMR) efflux transporter (qacG-like, with a secondary match to qacJ), and an aminoglycoside adenylyltransferase [ANT(9)-Ic] (*SI Table 5*). Screening with ResFinder further identified a class C (AmpC) β-lactamase, blaOCH-2–like, in both genomes (≈99.5% coverage, ≈83% identity). The high concordance between CARD and ResFinder results indicates a stable, efflux-dominated resistome complemented by enzymatic inactivation mechanisms against aminoglycosides and β-lactams. On the other hand, virulence factor screening using ABRicate (VFDB) detected only loci related to lipopolysaccharide (LPS) and cyclic β-1,2-glucan biosynthesis (kdsA/B, pgm, cgs, acpXL, manCoAg, manAoAg, wbpZ, htrB, fabZ) and the intracellular survival factor ricA. These are conserved structural or stress-response genes found across non-pathogenic *Brucella* /*Ochrobactrum* lineages and are more consistent with environmental adaptation than with host-specialized pathogenicity. No homologs of classical virulence determinants (toxins, adhesins, or secretion systems) were detected, supporting the classification of both isolates as low-pathogenic environmental variants. To validate the genomic resistome predictions, we performed phenotypic antimicrobial susceptibility testing (AST) of both landfill isolates using the VITEK 2 platform. The AST profiles were largely consistent with the genomic analysis (*SI Table 7*). These landfill isolates showed high resistance to piperacillin/tazobactam and intermediate susceptibility to several cephalosporins, including ceftriaxone, cefoperazone/sulbactam, and cefepime, consistent with the presence of the blaOCH-2–like AmpC β-lactamase identified in the genome. In contrast, landfill isolates remained susceptible to carbapenems (meropenem and imipenem), reflecting the absence of carbapenemase genes in the genomic screen. Additionally, susceptibility was observed for aminoglycosides (amikacin, gentamicin), fluoroquinolones (ciprofloxacin), and trimethoprim/sulfamethoxazole. The agreement between genotypic predictions and phenotypic AST results supports the presence of a limited intrinsic resistome primarily driven by efflux systems and β-lactamase activity, while the lack of major virulence determinants further indicates that these isolates represent environmental, low-pathogenic *Brucella intermedia* variants.

The observed resistance gene profile indicates that both *B. intermedia* isolates are environmentally resilient and well-adapted to the stress conditions commonly found in soil and landfill ecosystems. Dominant RND and SMR pumps indicate efflux-mediated detoxification that may extend beyond antibiotics to organic solvents and certain aromatic hydrocarbons that may be generated during plastic biodegradation. Before the widespread use of antibiotics, RND efflux pumps likely evolved to help bacteria detoxify naturally occurring compounds in their environment. The ability to pump out antibiotics is considered an evolutionary novelty that stems from their original function (Alvarez-Ortega et al., 2013). The presence of a chromosomal class C β-lactamase (blaOCH-2–like) is consistent with intrinsic β-lactam resistance typical of environmental *Brucella/Ochrobactrum* lineages (Li et al., 2021; Nadjar et al., 2001), suggesting a baseline tolerance to many penicillins and cephalosporins. This pattern likely reflects environmental adaptation rather than recent clinical antibiotic exposure. This is further supported by the virulence factor profiles of the two landfill isolates, as only conserved structural and stress-responsive virulence genes were detected. Loci involved in LPS and cyclic β-1,2-glucan biosynthesis and the intracellular survival factor ricA play roles in cell envelope integrity and osmotic stress tolerance rather than host infection (Byndloss and Tsolis, 2016). In addition, the complete absence of virulence-associated secretion systems, adhesins, or exotoxins clearly distinguishes these isolates from pathogenic *Brucella* strains (Byndloss and Tsolis, 2016). Taken together, the resistome, virulome, and phenotypic AST data support the view that isolates X and Y are environmentally resilient but low-pathogenic *B. intermedia* variants, consistent with adaptation to plastic-rich landfill habitats rather than host-associated pathogenic specialization.

### 3.6 LLDPE-degrading Brucella isolates form biofilms on plastic surfaces

Biofilm formation often helps bacteria persist under environmentally challenging conditions, particularly under nutrient limitation and during growth on hydrophobic substrates (Parrilli et al., 2022; Yin et al., 2019). To determine whether the landfill-derived *Brucella* isolates formed biofilms under plastic-associated conditions, we performed the Congo red agar (CRA) assay following Freeman et al. (1989) and additionally monitored surface-associated growth on LLDPE pieces (*SI Figure 6*). Under standard CRA conditions, neither isolate showed black pigmentation or precipitate formation, indicating no detectable biofilm-associated phenotype in the absence of the hydrocarbon supplement. In contrast, when we modified the medium with hexadecane, a simple polyethylene mimic (Gyung Yoon et al., 2012; Montazer et al., 2020), both isolates displayed black pigmentation and precipitates in both agar and broth cultures, consistent with biofilm-associated growth under hydrocarbon-substituted conditions.

The two isolates nevertheless differed in the spatial pattern and apparent strength of the biofilm phenotype. In the hexadecane-supplemented assay, isolate X showed robust growth and extensive black precipitation on both solid and liquid modified media, indicating strong biofilm-associated growth in the presence of hexadecane (*SI Figure 6A and 6B*). By contrast, isolate Y showed heterogeneous black precipitation on solid medium, with growth concentrated in regions where hexadecane droplets were visibly accumulated (*SI Figure 6C*). However, isolate Y also produced uniform black coloration in broth (*SI Figure 6D*), indicating that it retained substantial biofilm-forming capacity when hexadecane was more evenly available. These observations suggest that both isolates can form biofilms under hydrocarbon-associated conditions, but that isolate X expresses this phenotype more uniformly on solid medium.

Direct observation of LLDPE pieces further strengthened this interpretation. Over the incubation time course, both isolates produced visible surface-associated growth on LLDPE, supporting the conclusion that they can colonize the plastic surface itself (*SI Figure 6E–L*). Visual examination of the LLDPE films in the modified CRA system showed progressive colonization over 24–144 h. Initial attachment was observed mainly along the film edges at 24 h, followed by gradual expansion toward the central regions at 48 and 72 h, and by 144 h much of the plastic surface was covered by visible surface-associated growth. Notably, growth remained restricted to regions containing LLDPE, supporting the interpretation that the plastic itself served as the primary surface for attachment and colonization under these conditions. However, isolate X appeared to accumulate denser and more extensive surface-associated material than isolate Y, particularly at later time points, whereas isolate Y showed comparatively lighter and less continuous coverage. Thus, the LLDPE-surface assay was consistent with the hexadecane-based CRA results and supported the view that isolate X has a stronger overall biofilm-associated phenotype than isolate Y under the tested conditions.

Together, these observations indicate that both isolates favor biofilm-associated growth under hydrocarbon-associated, nutrient-limited conditions that more closely resemble plastic-rich environmental settings than conventional nutrient-replete media. The patchier phenotype of isolate Y on solid medium likely reflects the low solubility and uneven distribution of hexadecane in agar (*SI Figure 6C*), which would generate localized zones of substrate availability. Under this interpretation, the difference between the isolates is better viewed as variation in the spatial distribution or consistency of biofilm-associated growth, rather than as presence versus absence of that capacity. Consistent with this, both isolates formed visible surface-associated material on LLDPE and showed biofilm-associated pigmentation in broth (*SI Figure 6B,D*). SEM provided additional structural support for this interpretation (*Figure 2; SI Figure 4*). Bacteria-exposed LLDPE films showed surface irregularities together with adherent cells and localized clustered colonization for both isolates (*Figure 2B–C; SI Figure 4B–C*). In parallel, biomass recovered from broth cultures after incubation with LLDPE formed compact clump-like aggregates with matrix-like morphology for both isolates (*Figure 2D; SI Figure 4D*), indicating that LLDPE exposure was also associated with pronounced bacterial aggregation in the liquid phase. Similar surface-associated growth has been reported in hydrocarbon-degrading and plastic-associated bacteria, where biofilm formation can enhance persistence on hydrophobic substrates and improve cell–substrate interactions during degradation of poorly soluble compounds (Urbanek et al., 2018).

To explore whether this phenotype was accompanied by known biofilm-associated genetic features, we screened the genomes of the two isolates against a manually curated set of 63 candidate biofilm-related genes (SI Table 3). This analysis identified 13 and 12 significant homologs in *B. intermedia* isolates X and Y, respectively, corresponding to nine unique reference genes in the curated dataset. The two genomes showed highly similar homolog profiles, with most matches falling into functional categories related to carbohydrate/exopolysaccharide-associated processes and regulatory components. Candidate homologs included algD-like, pslA-like, pslB/AlgA-like, gacA-like, and related genes previously implicated in matrix formation or biofilm regulation in other bacterial systems. In *Pseudomonas*, the psl locus contributes to biofilm initiation and matrix formation, while alginate- and Psl-associated exopolysaccharides support biofilm architecture and persistence (Hentzer et al., 2001; Irie et al., 2012; Jackson et al., 2004). The close correspondence between isolates X and Y therefore suggests that both strains possess a broadly similar genomic repertoire of functions plausibly associated with biofilm-related surface growth.

For isolate X, transcriptomic data provided an additional line of support beyond its genome content. Under the LLDPE condition, isolate X expressed several homologs linked to carbohydrate-processing, exopolysaccharide-associated, and regulatory functions, including algD-like, pslA-like, pslB/AlgA-like, RhlR-like, and GacA-like candidates (*SI Figure 7*). Although homology-based annotation and condition-associated expression do not establish direct functional equivalence with the corresponding pathways described in model organisms such as *P. aeruginosa*, these data are consistent with the possibility that biofilm formation on LLDPE in isolate X is accompanied by expression of genes related to extracellular matrix-associated growth. In particular, the combined presence and expression of multiple carbohydrate-processing and putative exopolysaccharide-associated homologs support the interpretation that the observed phenotype reflects active surface-associated growth rather than passive material deposition alone.

Overall, our results indicate that the landfill-derived *B. intermedia* isolates possess a conserved set of candidate functions plausibly linked to biofilm formation and that both can establish biofilm-associated growth under plastic-relevant conditions. The more pronounced phenotype of isolate X in both surrogate and direct surface assays, together with LLDPE-associated expression of multiple exopolysaccharide-related and regulatory homologs, strengthens the inference that active matrix-associated growth accompanies surface colonization in *Brucella* landfill isolates. At the same time, the SEM data suggest that the observable architecture depends on the experimental context, ranging from adherent cells and localized clustered colonization on LLDPE surfaces to dense aggregate formation in recovered broth biomass. Although these data do not by themselves define the matrix polymer or assign isolate-level phenotypic differences to specific genes, they support a model in which biofilm formation contributes to plastic-associated persistence in these environmental isolates.

Taken together, the chemical, structural, genomic, transcriptomic, and biofilm-associated observations presented in Section 3 support a model of LLDPE biodegradation in the landfill-derived *B. intermedia* isolates, integrating surface colonization, oxidative modification, fragment uptake, and intracellular assimilation (*Figure 5*).

**Figure 5.**
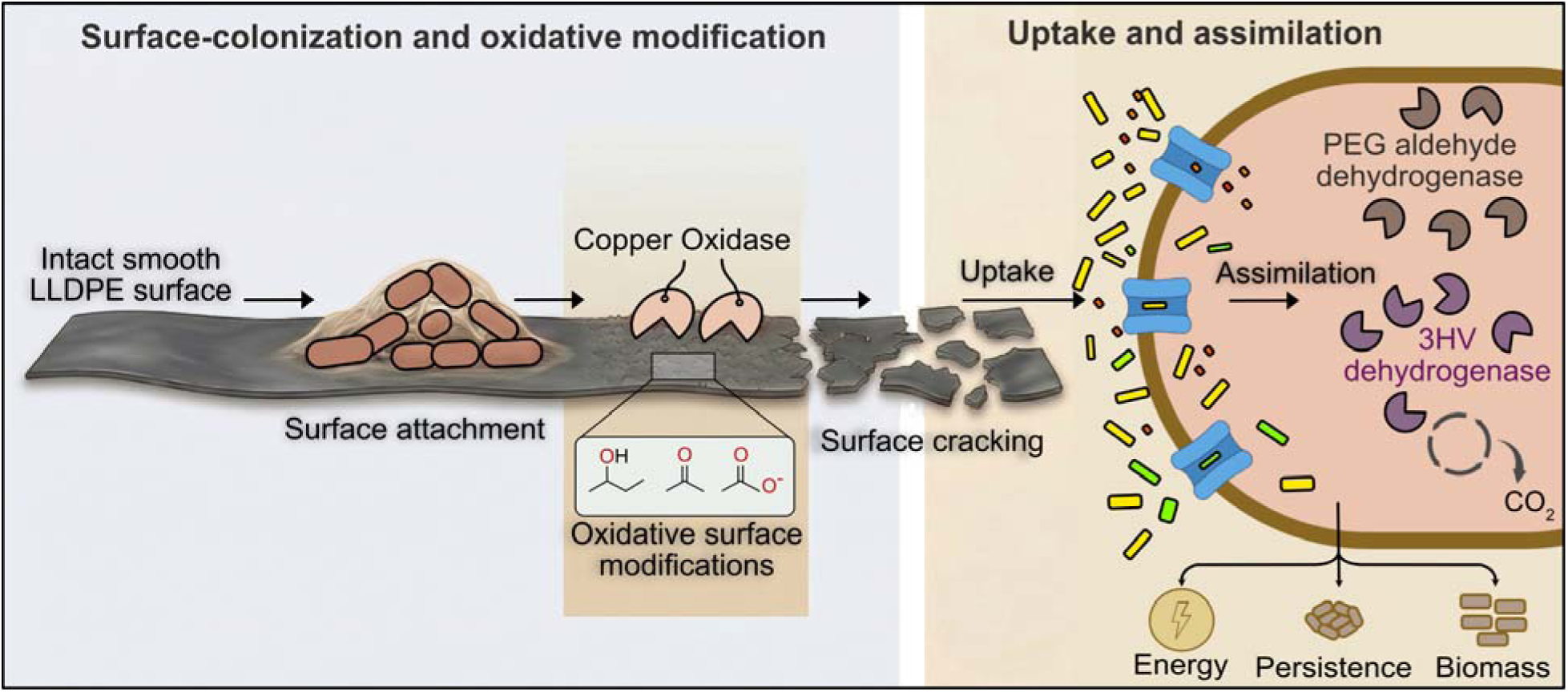
A model of LLDPE biodegradation in landfill-derived *Brucella intermedia*. The model summarizes the sequence of surface colonization and biofilm-associated growth on LLDPE, oxidative modification of the polymer surface, formation of an altered and roughened plastic substrate, uptake of small oxidized fragments, and intracellular assimilation. Together, these steps represent a conceptual framework for plastic-associated growth and biodegradation inferred from the combined functional, genomic, transcriptomic, and imaging data.

## 4. Conclusion

This study provides the first combined experimental, genomic, and transcriptomic evidence that landfill-derived *Brucella intermedia* isolates X and Y can initiate and sustain LLDPE biodegradation under carbon-limited conditions. Both isolates demonstrated the capacity to chemically and physically modify LLDPE, carried and expressed conserved candidate gene families spanning key biodegradation stages, and exhibited biofilm-associated growth under plastic-relevant conditions. At the same time, resistome, virulome, and AST analyses supported their classification as environmentally resilient, low-pathogenic variants. Together, these findings expand the known diversity of plastic-associated bacteria and position landfill-derived *B. intermedia* as an unexpected but ecologically relevant lineage for future studies of polymer degradation and plastic-associated microbial adaptation. More broadly, these findings argue for systematic microbiological exploration of plastic-rich niches as reservoirs of previously undescribed plastic-responsive and plastic-degrading microorganisms.

## Supporting information

Revised_Supplementary Information File 1

## CRediT authorship contribution statement

Ifthikhar Zaman: Writing – original draft, Writing – review & editing, Validation, Methodology, Investigation, Formal analysis, Data curation, Conceptualization, Visualization. Mahdi Muhammad Moosa: Writing – original draft, Writing – review & editing, Validation, Methodology, Investigation, Conceptualization, Visualization, Formal analysis, Data curation. M. Mahboob Hossain: Writing – review & editing, Validation, Supervision.

## Declaration of generative AI use

The author(s) used Grammarly and ChatGPT for language editing and ChatGPT, Gemini, and Grok for technical assistance in software setup and data analysis scripting. All content was subsequently reviewed and approved by the author(s), who assume full responsibility for the final manuscript.

## Data Availability

All sequencing data generated in this study are available in the ENA Project study accession PRJEB101032. Transcriptomic sequencing data and related analysis outputs are publicly available via Zenodo (<u>10.5281/zenodo.19098528</u>).

## Acknowledgments

The authors thank Zwad Al Saiyan for assistance with figure preparation, Maksudur Rahman Nayem for support with transcriptomic analysis, and Sadrina Afrin Mowna for proofreading the manuscript.

