## Supplementary material for "Functional, genomic, and transcriptomic insights into Linear Low-Density Polyethylene (LLDPE) biodegradation by landfill-derived *Brucella intermedia*": Revised_Supplementary Information File 1

| **Contents** | **Page** |
| --- | --- |
| SI Figure 1……………………………………………………………………………. | 2 |
| SI Figure 2……………………………………………………………………………. | 3 |
| SI Figure 3……………………………………………………………………………. | 4 |
| SI Figure 4……………………………………………………………………………. | 5 |
| SI Figure 5……………………………………………………………………………. | 6 |
| SI Figure 6……………………………………………………………………………. | 7 |
| SI Figure 7……………………………………………………………………………. | 9 |
| SI Table 1…………………………………………………………………………….. | 10 |
| SI Table 3…………………………………………………………………………….. | 12 |
| SI Table 7…………………………………………………………………………….. | 15 |
| SI Note 1……………………………………………………………………………… | 15 |
| Supplementary Information Table (provided as separate files) Legends…………….. | 17 |
| References…………………………………………………………………………….. | 18 |

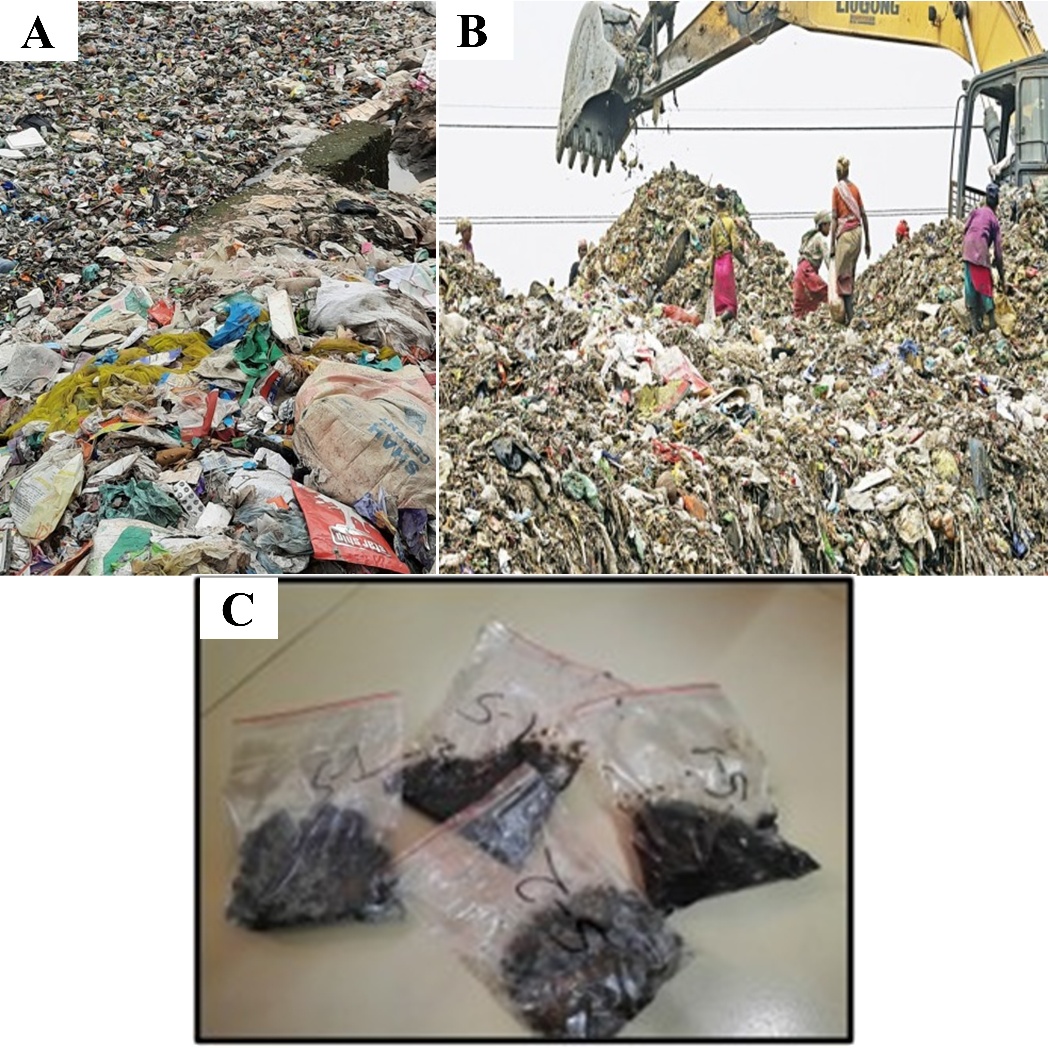

**SI Figure 1. Sampling site and soil sample collection at the Matuail landfill, Dhaka, Bangladesh. (A)** Drainage area clogged with mixed landfill waste, including plastic debris. **(B)** Ongoing waste disposal at the landfill site. **(C)** Collected soil samples stored in labeled plastic zipper bags.

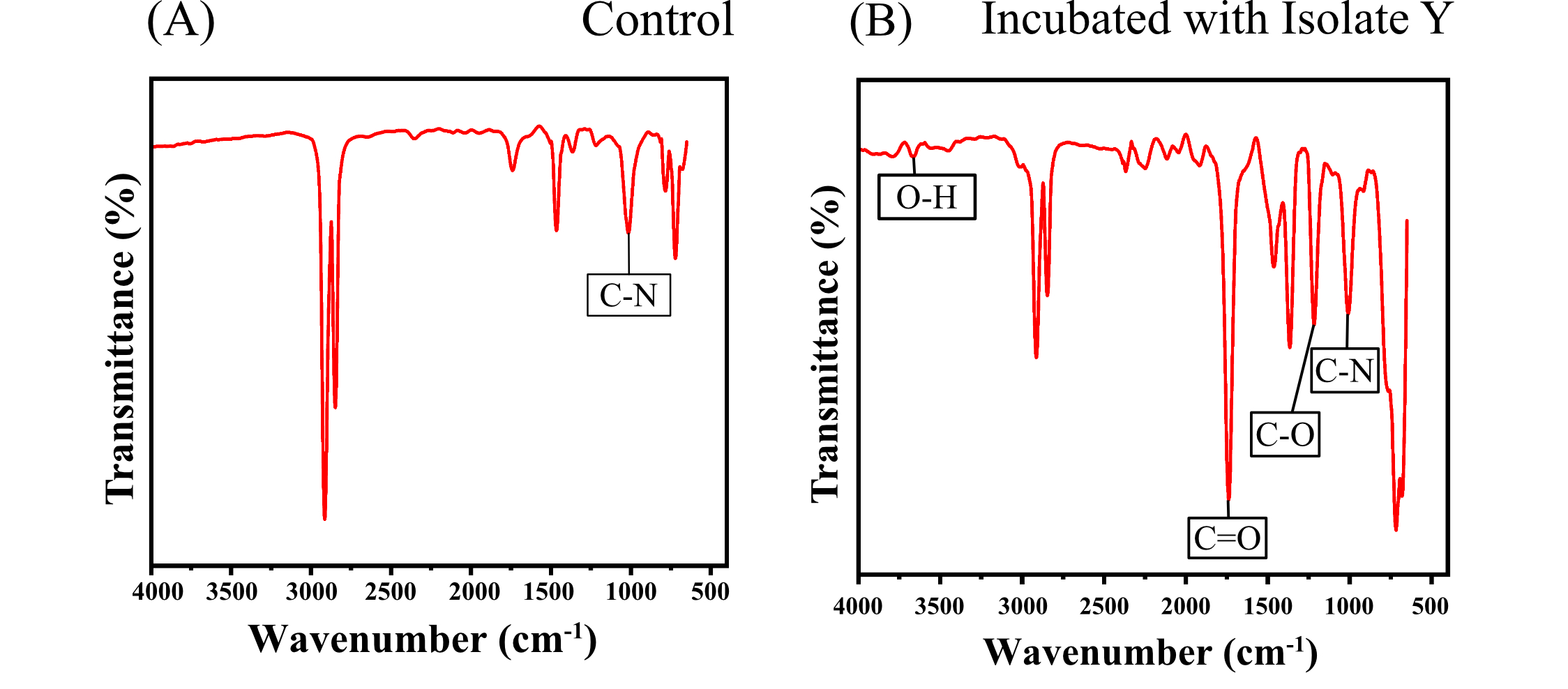

**SI Figure 2. FTIR spectra showing chemical modification of LLDPE following exposure to landfill *Brucella intermedia* isolate Y.** **(A)** FTIR spectrum of untreated control LLDPE, showing the characteristic polymer bands and a weak band near ~1020 cm⁻¹, possibly associated with additives or processing residues. **(B)** FTIR spectrum of LLDPE treated with isolate Y, showing additional bands assigned to O–H (3000–3670 cm⁻¹), C=O (1550–1720 cm⁻¹), and C–O (1250–1300 cm⁻¹) functional groups, consistent with oxidative modification of the polymer surface, similar to that observed for isolate X (see Figure 1, main text).

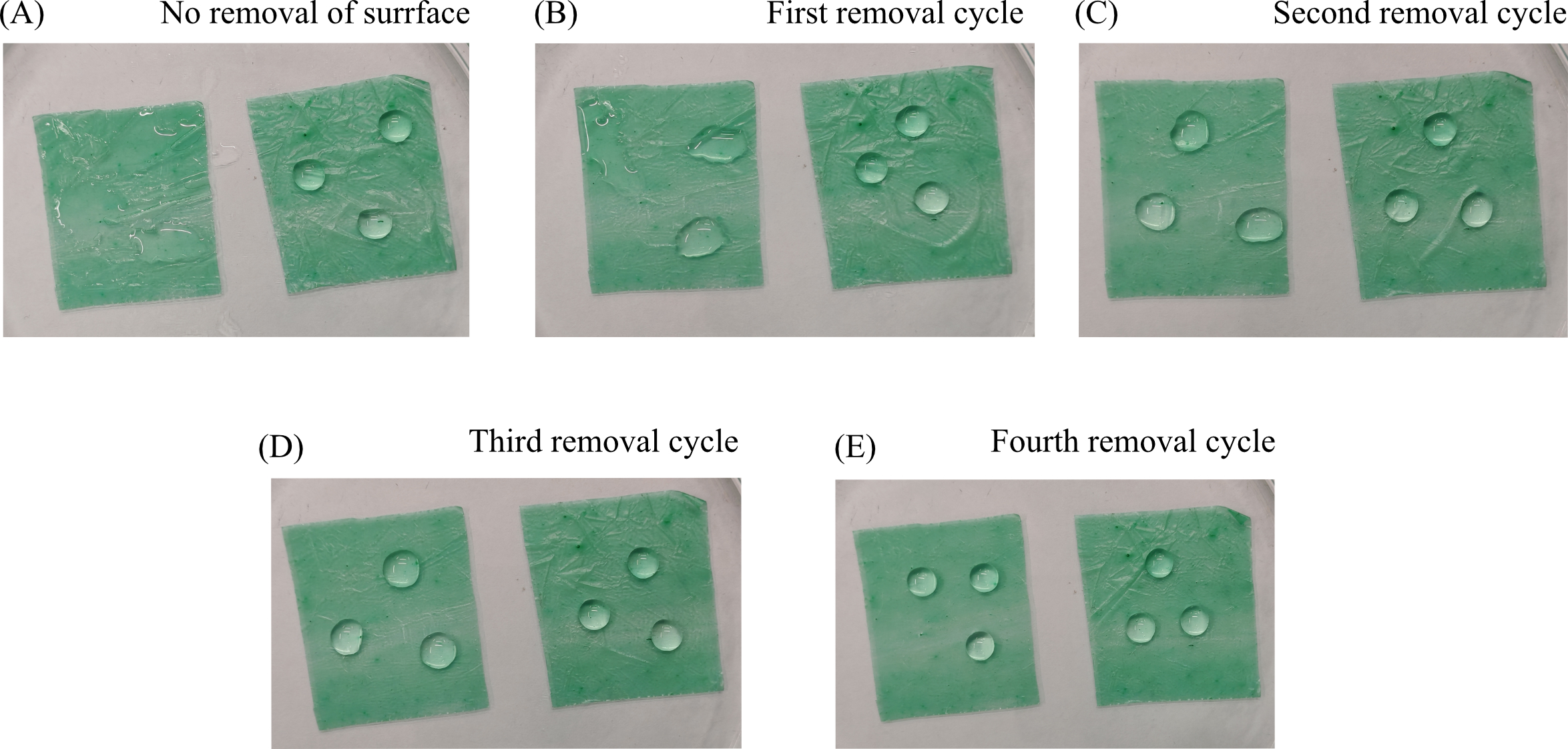

**SI Figure 3. Distilled water droplet spreading on control and isolate X-treated LLDPE across successive surface removal cycles.** In each panel, the left strip is isolate X-treated LLDPE and the right strip is an untreated control LLDPE. Three 100 µL distilled water droplets were placed on each strip, and droplet area was quantified using ImageJ. **(A)** Before surface removal, droplets on the treated LLDPE spread extensively and did not retain a well-defined spherical shape, whereas droplets on the control remained compact. This greater spreading is consistent with increased surface wettability of the treated LLDPE, likely reflecting oxidation-induced modification of the outermost polymer surface. **(B)** After the first surface removal cycle, two of the three droplets on the treated LLDPE began to retain a more defined shape, although droplet spreading remained greater than on the control. **(C)** After the second surface removal cycle, all three droplets on the treated LLDPE formed clearly defined droplets, but their contact area remained larger than that of the control. **(D)** After the third surface removal cycle, droplets on the treated LLDPE approached the size and shape observed on the control strip. **(E)** After the fourth surface removal cycle, treated and control LLDPE showed similar droplet sizes, indicating loss of the surface-associated wettability difference. Taken together, the progressive reduction in droplet spreading across successive surface removal cycles is consistent with the removal of an oxidized surface layer generated during exposure to isolate X, eventually revealing underlying material with wettability similar to that of the untreated control. Corresponding quantification is shown in the main text Figure 1C.

**SI Figure 4. Scanning electron micrograph analysis of LLDPE surface alterations and bacterial responses associated with LLDPE exposure in landfill *Brucella intermedia* isolate Y. (A)** Untreated control LLDPE showing a mostly smooth surface (same as Figure 2A, main text). **(B)** Cleaned LLDPE recovered after bacterial incubation with isolate Y, showing a roughened and irregular surface compared with the untreated control. **(C)** LLDPE surface with dense surface-associated bacterial growth and extensive coverage in the CRA system, suggestive of biofilm-like development. **(D)** Dense bacterial aggregates recovered by centrifugation from modified CRB cultures containing LLDPE, showing compact clump-like biomass with matrix-like morphology. Scale bars represent 3 µm.

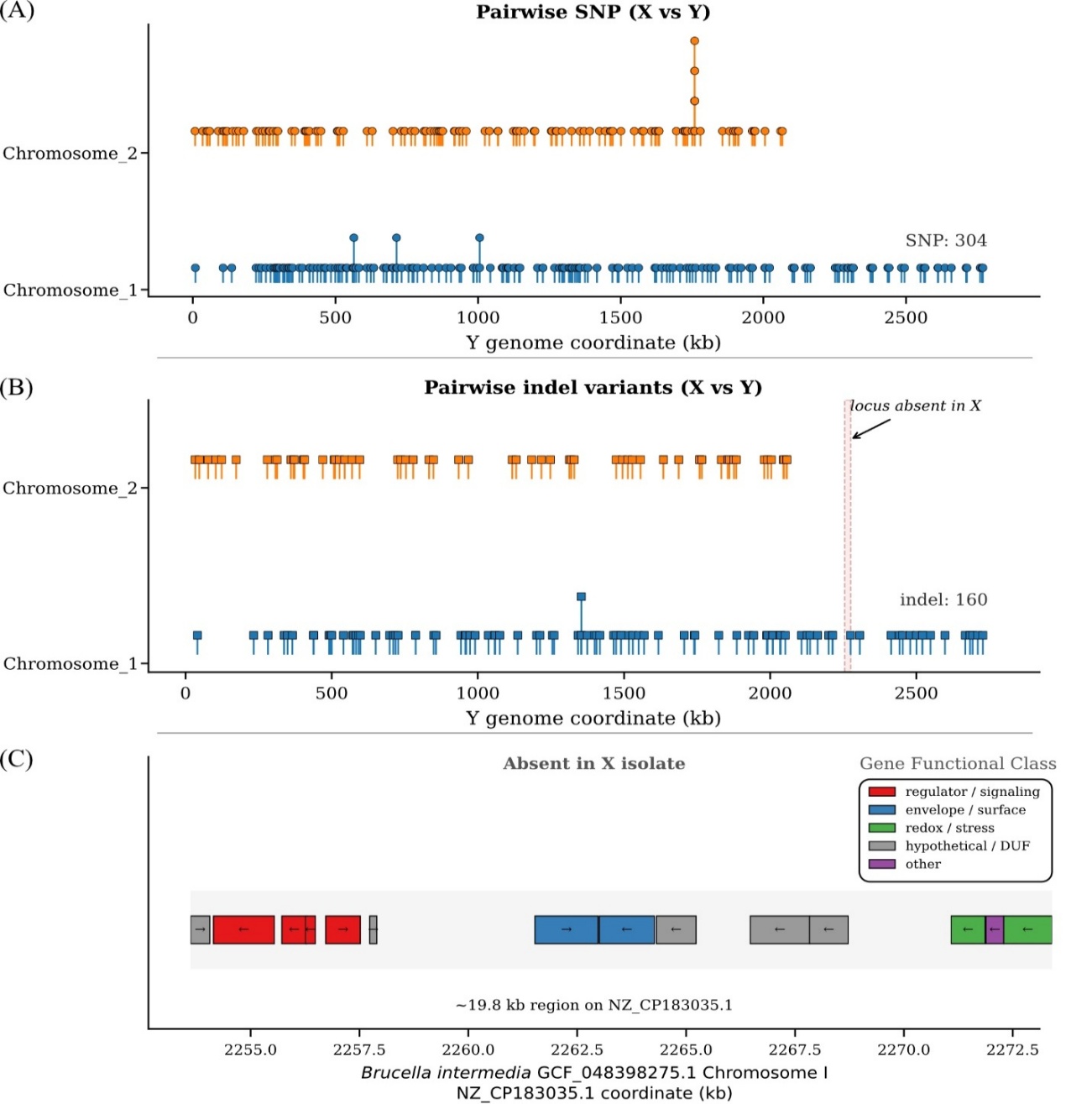

**SI Figure 5. Pairwise variant distribution and genomic differences between isolates X and Y.** **(A)** Distribution of single-nucleotide polymorphisms (SNPs) between isolates X and Y. Variants identified using Snippy with isolate Y as the reference are shown as lollipop markers across the two chromosomes; vertical stacking indicates closely spaced or overlapping variant positions. **(B)** Distribution of small insertion and deletion (indel) variants between isolates X and Y. Indels are distributed across both chromosomes, with an additional ~19.8 kb region on chromosome 1 absent in isolate X. **(C)** Gene content of the ~19.8 kb region absent in isolate X. Gene models were extracted from the annotation of the closest *Brucella intermedia* reference genome (GCF_043898275.1) and are colored by broad functional category, highlighting the loss of a contiguous locus in isolate X.

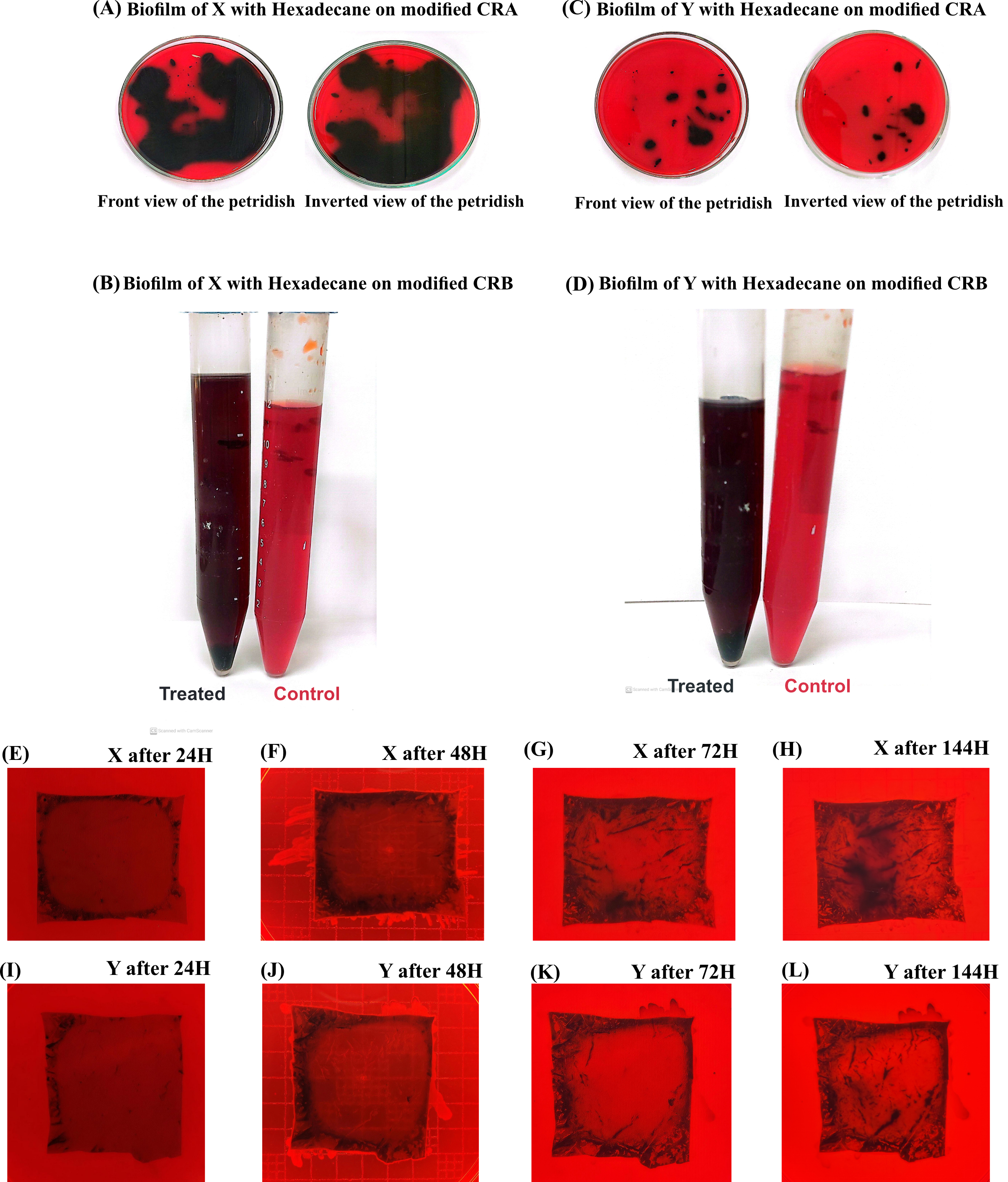

**SI Figure 6. Biofilm formation by landfill-derived isolates X and Y under hexadecane- and LLDPE-associated conditions.** Black pigmentation and precipitate formation in Congo red media indicate biofilm-associated growth. **(A)** Modified Congo red agar (CRA) supplemented with hexadecane showing biofilm formation by isolate X. **(B)** Modified Congo red broth supplemented with hexadecane showing biofilm formation by isolate X (left) and the negative control (right). **(C)** Modified CRA supplemented with hexadecane showing biofilm formation by isolate Y. **(D)** Modified Congo red broth supplemented with hexadecane showing biofilm formation by isolate Y (left) and the negative control (right). **(E–H)** Time-course of biofilm formation by isolate X on LLDPE in modified CRA medium at 24, 48, 72, and 144 h, respectively. Surface-associated growth was first visible mainly along the edges of the LLDPE film at 24 h and became progressively more extensive over time, with dense coverage evident by 144 h. **(I–L)** Time-course of biofilm formation by isolate Y on LLDPE in modified CRA medium at 24, 48, 72, and 144 h, respectively. Biofilm formation was also observed on the LLDPE surface, although coverage appeared less extensive than in isolate X over the same time course. In both isolates, visible growth was associated with the presence of the LLDPE film.

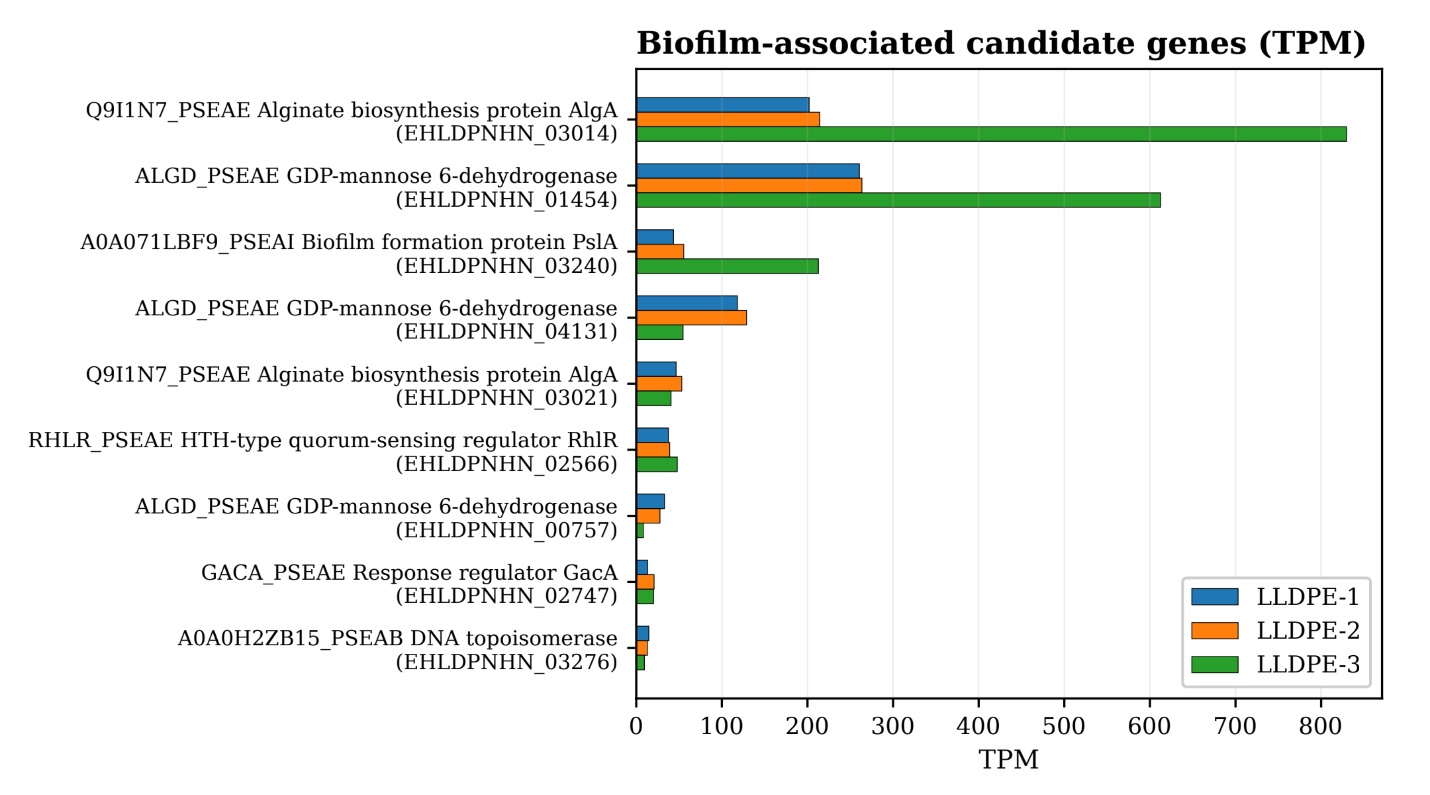

**SI Figure 7. Expression of candidate biofilm-associated homologs in *Brucella intermedia* isolate X under the LLDPE condition.** Horizontal bar plots show transcript abundance, reported as transcripts per million (TPM), for selected candidate homologs in three biological replicates grown under the LLDPE condition (LLDPE-1, LLDPE-2, and LLDPE-3). The y-axis shows the best-hit homolog annotation together with the corresponding isolate X locus tag (prokka annotation). Based on gene-name mapping of the best-hit annotations, the displayed candidates comprise three algD-like homologs (ALGD_PSEAE), two pslB-like homologs (Q9I1N7_PSEAE), and single pslA-like (A0A071LBF9_PSEAI), rhlR-like (RHLR_PSEAE), gacA-like (GACA_PSEAE), and pslN-like (A0A0H2ZB15_PSEAB) homologs. Multiple homologs linked to carbohydrate-processing, putative exopolysaccharide-associated, and regulatory functions were expressed, with algD-like and pslB-like homologs showing the highest transcript abundance among the candidates displayed.

**SI Table 1:** Representative *Brucella* species classified by ecological niche and pathogenic potential.

| **Name** | **Classification** | **Justification** | **Reference** |
| --- | --- | --- | --- |
| *Brucella pituitosa* | Environmental | Isolated from hydrocarbon-polluted sediment; environmental origin | (Mahjoubi et al., 2025) |
| *Brucella pseudogrignonensis* | Environmental/ Opportunistic | Emerging pathogen in patients, but derived from former *Ochrobactrum pseudogrignonense*; opportunistic | (Li et al., 2021) |
| *Brucella inopinata* | Intermediate/ Opportunistic | Atypical Brucella species with rare human cases | (Occhialini et al., 2022) |
| *Brucella ceti* | Intermediate/ Opportunistic | Marine mammal pathogen; zoonotic risk in handlers | (Nymo et al., 2011a) |
| *Brucella abortus* | Pathogenic | Classical zoonotic bovine/human brucellosis agent | (Qureshi et al., n.d.) |
| *Brucella melitensis* | Pathogenic | Most virulent Brucella species causing human brucellosis | (Qureshi et al., n.d.) |
| *Brucella pinnipedialis* | Intermediate/ Opportunistic | Seal/sea lion-associated Brucella | (Nymo et al., 2011b) |
| *Brucella ovis* | Animal pathogen | Sheep pathogen; non-zoonotic but pathogenic in livestock | (Poester et al., 2013) |
| *Brucella suis* | Pathogenic | Swine-associated brucellosis agent with human infection reports | (Poester et al., 2013) |
| *Brucella canis* | Intermediate/ Opportunistic | Dog pathogen with occasional human cases | (Qureshi et al., n.d.) |
| *Brucella microti* | Environmental | Isolated from voles and soil, an environmental reservoir | (Scholz et al., 2008) |
| *Brucella anthropi* | Environmental | Former *Ochrobactrum anthropi*; opportunistic soil/water isolate | (Sardana and Rajeev Verma, 2025) |
| *Brucella* sp. MAB 22 | Environmental | Environmental isolation from the cyanobacterial bloom lake | (“Brucella sp. MAB-22 genome assembly ASM2590847v1,” n.d.) |
| *Brucella pseudintermedia* | Environmental |  | (Yang et al., 2024) |
| *Brucella intermedia* | Environmental | Former *Ochrobactrum intermedium*; hospital/environmental strains | (Yang et al., 2024) |

**SI Table 3:** Curated list of biofilm formation/regulation genes from different species.

| **Organism** | **Gene(s)** | **Notes / Function** | **References** |
| --- | --- | --- | --- |
| ***Staphylococcus aureus*** | **icaA / icaD** | Part of the *icaADBC* operon responsible for PIA (polysaccharide intercellular adhesin) synthesis, key for biofilm matrix. | (Ahmad et al., 2022) |
|  | **bap** | Biofilm-associated protein that mediates cell–surface and cell–cell adhesion. | (Arciola et al., 2015) |
|  | **sarA** | Global regulator enhancing biofilm formation via ica expression and surface protein regulation. | (Tormo et al., 2005) |
|  | **agrA / agrC** | Quorum-sensing regulator; controls biofilm dispersal; *agr* mutants often form stronger biofilms. | (Juszczuk-Kubiak, 2024) |
|  | **fnbA / fnbB** | Fibronectin-binding proteins that mediate initial adhesion to host extracellular matrix. | (Cegelski et al., 2009) |
| ***Pseudomonas aeruginosa*** | **pelA–pelF** | Involved in Pel polysaccharide biosynthesis, an essential matrix component. | (Van Loon et al., 2025) |
|  | **pslA–pslN** | Synthesizes PSL exopolysaccharide, critical for biofilm initiation and structure. | (Yu et al., 2016) |
|  | **algD** | Encodes GDP-mannose dehydrogenase, key enzyme in alginate biosynthesis for mucoid biofilms. | (Hulen, 2023) |
|  | **lasI / lasR** | Part of quorum-sensing (Las system); regulates biofilm maturation and virulence genes. | (Schlichter Kadosh et al., 2024) |
|  | **rhlI / rhlR** | Rhl quorum-sensing system regulates rhamnolipid and other biofilm-related factors. | (Mukherjee et al., 2017) |
|  | **gacA / gacS** | Two-component regulatory system controlling biofilm genes and exopolysaccharide production. | (Song et al., 2023) |
| ***Escherichia coli*** | **csgA / csgB** | Curli fimbriae subunits are responsible for cell adhesion and surface attachment. | (Swasthi and Mukhopadhyay, 2017) |
|  | **bssS / bssR** | Regulate biofilm formation and motility via stress response pathways. | (Domka et al., 2006) |
|  | **fimH** | Encodes adhesin at the tip of type 1 fimbriae; critical for initial attachment. | (Sauer et al., 2016) |
| ***Vibrio cholerae*** | **luxS** | Quorum-sensing gene involved in AI-2 synthesis; modulates biofilm formation. | (Wang et al., 2019) |
|  | **vpsA–vpsL** | Vibrio polysaccharide biosynthesis locus, essential for matrix formation. | (Gao et al., 2020) |
|  | **hapR** | Negative regulator of biofilm formation; hapR mutants form robust biofilms. | (Caigoy et al., 2022) |
| ***Bacillus subtilis*** | **epsE** | Encodes a glycosyltransferase for EPS synthesis in the biofilm matrix. | (Guttenplan et al., 2010) |
|  | **tapA / tasA** | Structural components of amyloid fibers in the biofilm matrix. | (Romero et al., 2011) |
|  | **sinR / sinI** | Master regulators controlling matrix gene expression, *sinR* represses, *sinI* derepresses biofilm genes. | (Dannenberg et al., 2023) |
| ***Enterococcus faecalis*** | **epa locus** | Encodes Enterococcal polysaccharide antigen important for biofilm formation and immune evasion. | (Norwood et al., 2024) |
|  | **gelE** | Gelatinase contributes to biofilm structure and virulence. | (Pirbonyeh et al., 2024) |
| ***Candida albicans*** | **bcr1** | Transcription factors required for adhesion and biofilm development. | (Nobile et al., 2006) |
|  | **efg1** | Regulates morphogenesis and biofilm formation. | (Connolly et al., 2013) |
|  | **hwp1** | Hyphal wall protein is essential for adhesion to host surfaces and biofilm integrity. | (Nobile et al., 2006) |

**SI Table 7:** Phenotypic antimicrobial susceptibility testing (AST) performed using the VITEK 2 platform, showing resistance (R), intermediate resistance (I), and susceptibility (S) of different antibiotics, where both isolates showed a similar resistance/susceptibility pattern.

| **Sl** | **Antibiotic** | **Class** | **MIC Value** | **Interpretation** |
| --- | --- | --- | --- | --- |
| 1 | Piperacillin/Tazobactam | Ureidopenicillin + β-lactamase inhibitor (β-lactam) | >= 128 | R |
| 2 | Ceftriaxone | 3rd-generation cephalosporin (β-lactam) | 32 | I |
| 3 | Cefoperazone/Sulbactam | 3rd-generation cephalosporin + β-lactamase inhibitor | 32 | I |
| 4 | Cefepime | 4th-generation cephalosporin (β-lactam) | 16 | I |
| 5 | Imipenem | Carbapenem (β-lactam) | 2 | S |
| 6 | Meropenem | Carbapenem (β-lactam) | 0.5 | S |
| 7 | Amikacin | Aminoglycoside | 8 | S |
| 8 | Gentamicin | Aminoglycoside | 4 | S |
| 9 | Ciprofloxacin | Fluoroquinolone | 0.5 | S |
| 10 | Trimethoprim/Sulfamethoxazole | Folate synthesis inhibitors | <= 20 | S |

**SI Note 1: Definition and interpretation of closest *Brucella intermedia* (****GCF_048398275.1)-specific accessory genomic islands.**

To identify genomic regions present in the hospital-wastewater *Brucella intermedia* reference strain but absent from our landfill isolates (X and Y), we aligned each isolate to the reference assembly and quantified per-base coverage for each annotated coding sequence (CDS). A CDS was considered “present at the locus” if ≥80% of its length showed nonzero coverage in that isolate. CDS that failed this coverage criterion were then searched against the isolate’s own assembly; if we detected a relocated match covering ≥80% of the CDS sequence with ≥90% amino acid identity, the CDS was classified as “present moved.” CDS that were neither present at the locus nor present moved were designated “absent.”

We then grouped absent CDS into “accessory islands” by merging CDS that lie on the same reference contig and are separated by ≤5 kb. For each resulting island, we report the reference coordinates, total span, number of CDS, GC content across the span, and representative annotated products (“example products”). This procedure minimizes noise from single borderline ORFs and instead captures coherent mobile/stress islands. A summary of these analyses is provided in the SI Table 2.

Most islands missing from both X and Y (“absent both” in the loss pattern column of the SI Table 2) were enriched for phage- and plasmid-associated functions, including IS3/IS5-family transposases, integrases, partition/replication factors (RepB/ParB/Spo0J), restriction–modification enzymes, DNA adenine methylases, lytic amidases, terminase subunits, portal proteins, and tail fiber/tape measure proteins. These islands had a lower GC content *vs.* genomic backgrounds as evidenced by dips in the circular plot (2^nd^ co-centric circle from the center, main-text Figure 3B). The observed gene content and atypical GC composition of these islands indicate that these genomic segments are *horizontally acquired accessory DNA*, perhaps related to hospital wastewater adaptation of the reference isolate, GCF_048398275.

We also identified large, atypically high-GC (~0.63) islands encoding multiple regulatory and oxidative stress–associated functions, including MarR- and LysR-family transcriptional regulators and peptide-methionine (R)-S-oxide reductase MsrB, as well as uncharacterized DUF-rich loci and a recombinase-like protein (SI Table 2). The combination of high GC content, stress-response/repair enzymes (*e.g.*, MsrB), and recombination-associated genes suggests that this block represents horizontally acquired stress-adaptation cargo present in the hospital-wastewater reference strain but absent from both landfill isolates.

These loci likely reflect niche-specific stress and mobile cargo in the hospital-wastewater isolate rather than core *B. intermedia* metabolism.

In contrast, smaller islands absent only from isolate X (“absent_X_only” in the loss_pattern column of the SI Table 2) encode a predicted two-component regulatory module (sensor histidine kinase plus an alternative sigma factor) and glycosyltransferase/oligosaccharide flippase/oxidoreductase enzymes involved in surface or envelope modification and redox tailoring. Isolate Y retains these loci. Together with the measured SNP/indel divergence between X and Y, this supports that X and Y are closely related but genomically distinct strains, rather than clonal replicates.

Notably, none of the missing islands encode canonical *Brucella* virulence determinants (e.g. type IV secretion systems, adhesins, classical toxins). This, along with the resistome/virulome profiling of X and Y, supports that these landfill isolates represent low-pathogenic, environmentally adapted *B. intermedia* strains.

**Supplementary Information Table (provided as separate files) Legends**

**SI Table 2.** Accessory genomic islands present in the hospital-wastewater *Brucella intermedia* (GCF_048398275.1) but are missing from landfill isolates X and/or Y.
loss_pattern: “absent_both” = island missing in both X and Y; “absent_X_only” = present in Y but missing in X. Coordinates are given relative to the reference genome (GCF_048398275.1). example_products lists representative annotated CDS within each island.

**SI Table 4:** Distribution of putative plastic-degrading enzymes across sequenced *Brucella* genomes.

**SI Table 5:** Predicted antimicrobial resistance genes identified in the landfill *Brucella* isolates X and Y.

**SI Table 6:** Summary of whole-genome average nucleotide identity (ANI) results for *Brucella* isolates X and Y, compared against *Brucella* reference genomes. ANI results were generated using FastANI, comparing the query genomes *vs.* all complete *Brucella* genomes available at the NCBI genomes database.
